# Uridine Metabolism as a Targetable Metabolic Achilles’ Heel for chemo-resistant B-ALL

**DOI:** 10.1101/2025.01.27.635108

**Authors:** Yuxuan Liu, Haowen Jiang, Jingjing Liu, Lucille Stuani, Milton Merchant, Astraea Jager, Abhishek Koladiya, Ti-Cheng Chang, Pablo Domizi, Jolanda Sarno, Tim Keyes, Dorra Jedoui, Ao Wang, Jodie Meng, Felix Hartmann, Sean C. Bendall, Min Huang, Norman J. Lacayo, Kathleen M. Sakamoto, Charles G. Mullighan, Mignon Loh, Jiyang Yu, Jun Yang, Jiangbin Ye, Kara L. Davis

## Abstract

Relapse continues to limit survival for patients with B-cell acute lymphoblastic leukemia (B-ALL). Previous studies have independently implicated activation of B-cell developmental signaling pathways and increased glucose consumption with chemo-resistance and relapse risk. Here, we connect these observations, demonstrating that B-ALL cells with active signaling, defined by high expression of phosphorylated ribosomal protein S6 (“pS6+ cells”), are metabolically unique and glucose dependent. Isotope tracing and metabolic flux analysis confirm that pS6+ cells are highly glycolytic and notably sensitive to glucose deprivation, relying on glucose for *de novo* nucleotide synthesis. Uridine, but not purine or pyrimidine, rescues pS6+ cells from glucose deprivation, highlighting uridine is essential for their survival. Active signaling in pS6+ cells drives uridine production through activating phosphorylation of carbamoyl phosphate synthetase (CAD), the enzyme catalyzing the initial steps of uridine synthesis. Inhibition of signaling abolishes glucose dependency and CAD phosphorylation in pS6+ cells. Primary pS6+ cells demonstrate high expression of uridine synthesis proteins, including dihydroorotate dehydrogenase (DHODH), the rate-limiting catalyst of *de novo* uridine synthesis. Gene expression demonstrates that increased expression of *DHODH* is associated with relapse and inferior event-free survival after chemotherapy. Further, the majority of B-ALL genomic subtypes demonstrate activity of DHODH. Inhibiting DHODH using BAY2402232 effectively kills pS6+ cells *in vitro*, with its IC50 correlated with the strength of pS6 signaling across 14 B-ALL cell lines and patient-derived xenografts (PDX). *In vivo* DHODH inhibition prolongs survival and decreases leukemia burden in pS6+ B-ALL cell line and PDX models. These findings link active signaling to uridine dependency in B-ALL cells and an associated risk of relapse. Targeting uridine synthesis through DHODH inhibition offers a promising therapeutic strategy for chemo-resistant B-ALL as a novel therapeutic approach for resistant disease.

## Main

Metabolic dysregulation is a hallmark of cancer with roles in tumor initiation, progression, and chemotherapy resistance^1–3^. Warburg first described cancer cells favoring glucose metabolism by glycolysis to produce lactate over oxidative phosphorylation even in the presence of sufficient oxygen^4^. Since this seminal observation, the understanding of cancer-specific metabolic adaptations has led to novel understanding of cancer biology and therapeutic opportunities. For example, both acute myeloid leukemia and glioma are characterized by mutations in isocitrate dehydrogenase (IDH), now a prominent therapeutic target^5–9^. Despite the recognition of dysregulated metabolism in cancer at large, its role in acute lymphoblastic leukemia (ALL) remains incompletely understood.

ALL is a cancer comprised of immature lymphocytes, most commonly of the B-lineage. ALL primarily affects children and adolescents as the most common malignancy in this age group, while approximately 20% of adult acute leukemia is ALL^10–12^. Prognosis is related to several factors including underlying genomic subtype, with the incidence of poor prognosis features and risk of relapse increasing with age such that teenagers and adults are more likely to have ALL with high-risk features and suffer relapse. In an effort to better understand ALL cells capable of mediating relapse, we identified a subset of pre-B-like ALL cells characterized by phosphorylation of several proteins: ribosomal protein S6 (pS6), 4EBP1, CREB, SYK, collectively termed “pS6+ cells”, whose presence at diagnosis was highly predictive of relapse^13^. Active signaling in B-cell developmental pathways, including IL-7 receptor (IL-7R) and pre-B cell receptor (pre-BCR) pathways, has previously been implicated in the pathogenesis and prognosis of ALL^14–17^. However, therapeutic targeting of this active signaling has not yet resulted in improved outcomes for these patients.

In the present study, we demonstrate that pS6+ cells possess a unique metabolic dependency on glucose. pS6+ cells utilize glucose carbons along the pentose phosphate and *de novo* pyrimidine synthesis pathways to produce uridine. We find that the dependency on glucose is directly correlated with the strength of pS6 signaling which is reversed when the signaling is inhibited. pS6+ cells drive uridine synthesis through phosphorylation of CAD, the enzyme catalyzing the initial steps of uridine synthesis. Evaluation of gene expression data from primary B-ALL patients demonstrates that higher expression of uridine synthesis genes is associated with relapse and worse event-free survival, consistent with our published observation relating these cells with relapse. Finally, targeting DHODH, the rate-limiting enzyme in *de novo* uridine synthesis, demonstrates remarkable efficacy in pS6+ B-ALL models across genetic backgrounds, including those at high relapse risk. Together, these data provide compelling evidence for a novel metabolic intervention relevant to patients at high risk of relapse with B-ALL.

## Results

### pS6+ cells have distinct metabolic gene signatures and energetics

Previously, we identified pS6+ cells in B-ALL patients at the time of diagnosis and found them to be associated with future relapse after standard chemotherapy^13^. To identify distinguishing features of relapse-associated pS6+ cells compared to pS6-cells from patients in continuous remission, we sorted pre-B cells from six diagnostic patient bone marrow (BM) samples with known pS6 status from our previously published B-ALL cohort (n = 3 pS6+; n = 3 pS6-; Supplemental Table 1 and 2) and performed whole transcriptome sequencing (Fig. 1A and Extended Data Fig. 1A and 1B). Pathway analysis demonstrated enrichment in mTORC1 and PI3K/Akt/mTOR signaling pathways in pS6+ cells, consistent with our published proteomic signature^13^. Additionally, pS6+ cells have higher expression of MYC gene targets and genes related to several metabolic pathways, including oxidative phosphorylation (OXPHOS), glycolysis, and fatty acid metabolism (Fig. 1B, Supplemental Table 3). These data confirmed the activation of PI3K/mTOR signaling we previously observed by proteomic analysis and suggested this signaling indicates a unique metabolic state.

**Fig. 1:**
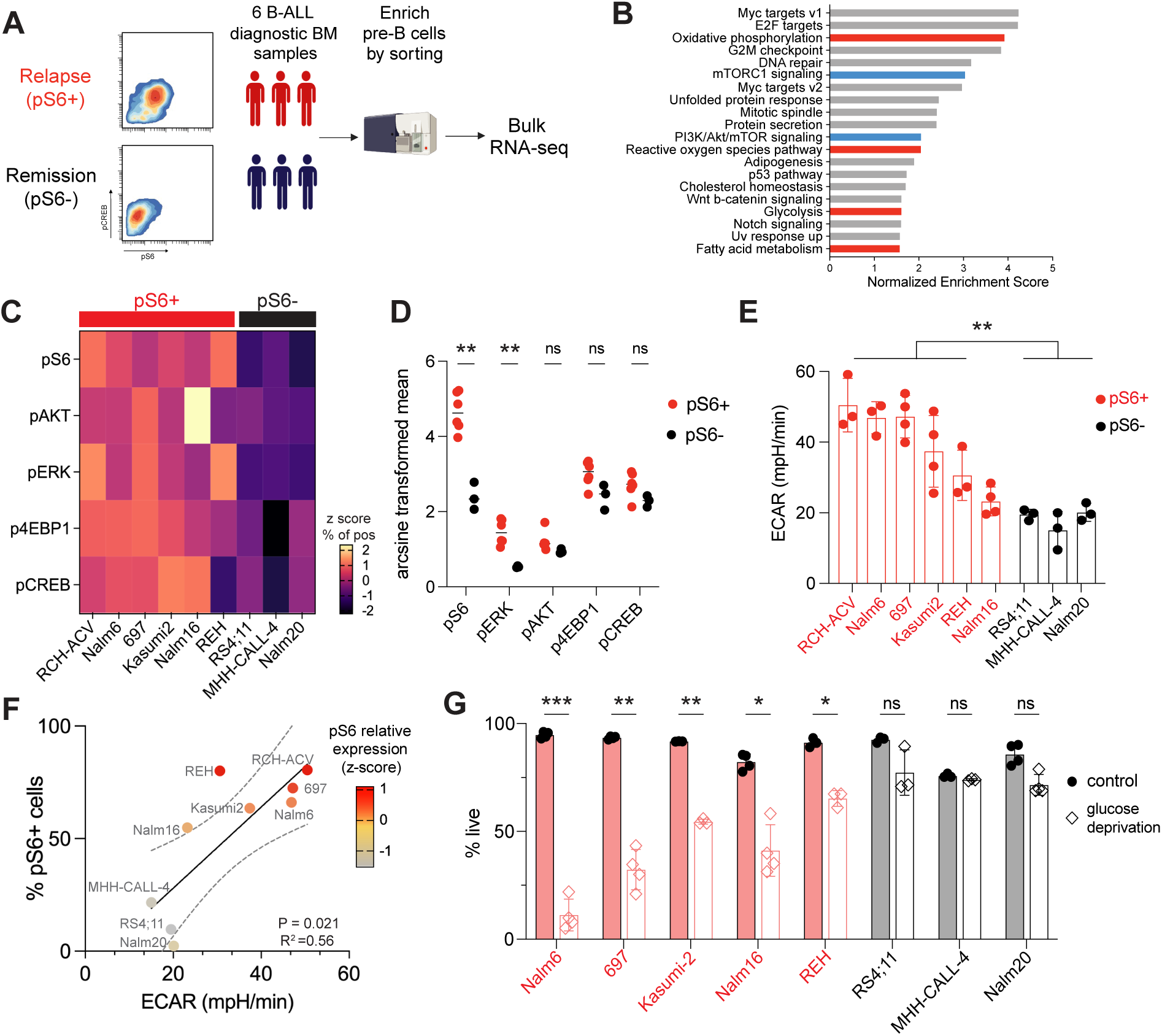
pS6+ cells have distinct metabolic gene signatures and are glucose dependent. **A.** Whole transcriptome sequencing was performed in primary diagnostic bone marrow (BM) samples from known pS6+ patients who would go on relapse (n=3) and pS6-patients who remain in continued remission (n=3). **B.** Differential expression analysis between primary pS6+ and pS6-cells. Gene set enrichment analysis (GSEA) was performed with the Hallmark database (FDR < 0.05). Diagnostic pS6+ cells are enriched for genes in PI3K and mTOR pathways (blue) as well as several metabolic pathways (red). **C.** Z-score based on frequency of cells positive for phosphorylated S6, AKT, ERK, 4EBP1 and CREB in B-ALL cell lines in basal state defines pS6+ lines (n=6, Nalm6, RCH-ACV, 697, Kasumi2, Nalm16 and REH) and pS6-lines (n=3, RS4;11, Nalm20, MHH-CALL-4). **D.** Expression (arcsinh transformed mean) of pS6 (S235/236), pAKT (S273), pERK (T202/Y204) and pCREB (S133) in pS6+ and pS6-cell lines (pS6, P = 0.002; pERK, P=0.004; pAKT, p= 0.095, ns; p4EBP1, p=0.095, ns; pCREB, p = 0.095, ns). **E.** Extracellular acidification rate (ECAR) indicating glycolytic activity in pS6+ cell lines (red) compared to pS6-cell lines (black; p = 0.0039). **F.** Correlation between the frequency of pS6+ cells and the glycolytic activity (measured by ECAR) in B-ALL cell lines (n=9, p = 0.021, R^2^ = 0.56). Each dot represents individual cell line colored by pS6 relative expression level (z-score of arcsinh transformed mean value) measured by mass cytometry. **G.** Cell viability after culture in medium with or without glucose (open bar) for 48 hours in pS6+ cells (red, n = 5) and pS6-cells (gray, n = 3). Cell death is measured by annexin V and PI staining by flow cytometry. Nalm6 p = 0.00098; 697 p = 0.0064; Kasumi2 p = 0.0022; Nalm16 p = 0.026; REH p = 0.0183; RS4;11, p = 0.614; MHH-CALL-4 p = 0.523; Nalm20 p = 0.081. Three or four biological replicates of experiments were performed. All data in dot plots and bar graphs are mean ± SD. Statistical tests used were Welch’s t test followed by Holm-Sidak multiple comparison test (D); Welch’s t test (E, I); and multiple paired t test followed using Šídák-Bonferroni method (G). ns, not significant, *p < 0.05, **p < 0.01, ***p < 0.001.

Using mass cytometry (CyTOF), we assessed the signaling status of nine B-ALL cell lines and categorized them into pS6+ or pS6-groups based on the frequency and signaling strength of pS6+ cells (Fig. 1C). Six cell lines are pS6+ (Nalm6, 697, RCH-ACV, Kasumi2, Nalm16, and REH), while three cell lines are pS6-(Nalm20, RS4;11, and MHH-CALL-4). In addition to pS6 activation (mean 4.7 *vs.* 2.4 p = 0.002), pS6+ cells have higher expression of pERK (1.5 *vs.* 0.5; p = 0.004), pAKT (1.2 *vs.* 0.9, p = 0.095, ns), p4EBP1 (3.0 *vs.* 2.4, p = 0.095, ns) and pCREB (p = 2.7 *vs.* 2.3, p = 0.095, ns, Fig. 1D). These results are in line with our previous observation in primary patient samples^13^.

To evaluate if the active signaling in pS6+ cells is associated with differences in metabolism, we performed metabolic flux assays in the B-ALL cell lines using Seahorse. Compared to pS6-cells, pS6+ cells are more energetic with significantly higher glycolysis and OXPHOS activity as indicated by basal extracellular acidification rate (ECAR, p = 0.0039) and oxygen consumption rate (OCR, p = 0.0191; Fig. 1E and Extended Data Fig. 2A). Further, both ECAR and OCR correlated with the frequency of pS6+ cells in B-ALL cell lines (ECAR, R^2^= 0.56, p = 0.021, Fig. 1F; and OCR, R^2^ = 0.67, p = 0.0068, Extended Data Fig. 2B). Thus, as suggested by the transcriptomic analysis of primary pS6+ cells, pS6+ cells have higher metabolic activity than pS6-cells, which is directly correlated to the frequency of pS6+ cells.

### pS6+ cells utilize glucose to fuel uridine synthesis

Glucose and glutamine serve as the primary carbon sources for glycolysis and OXPHOS in cancer cells^18^. To uncover the metabolic dependencies of pS6+ cells, we cultured B-ALL cell lines under glucose or glutamine deprivation conditions. pS6+ cells were dependent on glucose for survival, with significant cell death occurring after 48 hours in a glucose-deprived medium (Nalm6 p = 0.00098; 697 p = 0.0064; Kasumi2 p = 0.0022; Nalm16 p = 0.026; REH p = 0.018; Fig. 1G). By contrast, pS6-cells were tolerant to glucose deprivation (RS4;11 p = 0.61; MHH-CALL-4 p = 0.52; Nalm20 p = 0.08, Fig. 1G). Glutamine deprivation did not increase cell death in any of the cell lines except for Nalm6 (Extended Data Fig. 2C).

To understand how pS6+ cells utilize glucose, we performed isotype tracing with U-^13^C-glucose in the absence of glutamine in pS6+ and pS6-cells. pS6+ cells distinctly incorporated ^13^C-glucose into metabolites in the glycolysis, pentose phosphate pathway (PPP), and the TCA cycle as illustrated schematically in Fig. 2A. We did not observe significant differences in fractional ^13^C labeling of intermediates in glycolysis and TCA cycle between pS6+ and pS6-cells (Extended Data Fig. 3). The labelling of individual intermediates is shown in Extended Data Fig. 4. In particular, pS6+ cells had significantly higher fractional ^13^C labeling in m+5 UDP (p = 0.0233), UTP (p = 0.002), and ATP (p = 0.0342) compared to pS6-cells (Fig. 2B), suggesting pS6+ cells are using glucose for PPP and nucleotide production. To understand what metabolites downstream of PPP and nucleotide production are essential for survival in pS6+ cells, we performed a rescue experiment where we cultured B-ALL cell lines in a glucose-deprived condition and evaluated cell survival after supplementing different metabolites (Fig. 2C). Uridine, as the first product of pyrimidine synthesis, most effectively rescued pS6+ cells from glucose deprivation (Nalm6, p < 0.0001; 697, p < 0.0001; Nalm16, p = 0.0002; and REH, p = 0.0485) while, as expected, there was no impact in pS6-cells. Pyruvate provided partial rescue against cell death in two cell lines (697, p = 0.0007; Nalm16, p = 0.016), indicating it can support glycolysis and TCA cycle flux as a key metabolite but is not sufficient for complete metabolic compensation. Other metabolites, including aspartate, pyrimidine and purine failed to rescue cells from death, highlighting glucose-dependent uridine synthesis as a critical vulnerability in pS6+ B-ALL cells.

**Fig. 2:**
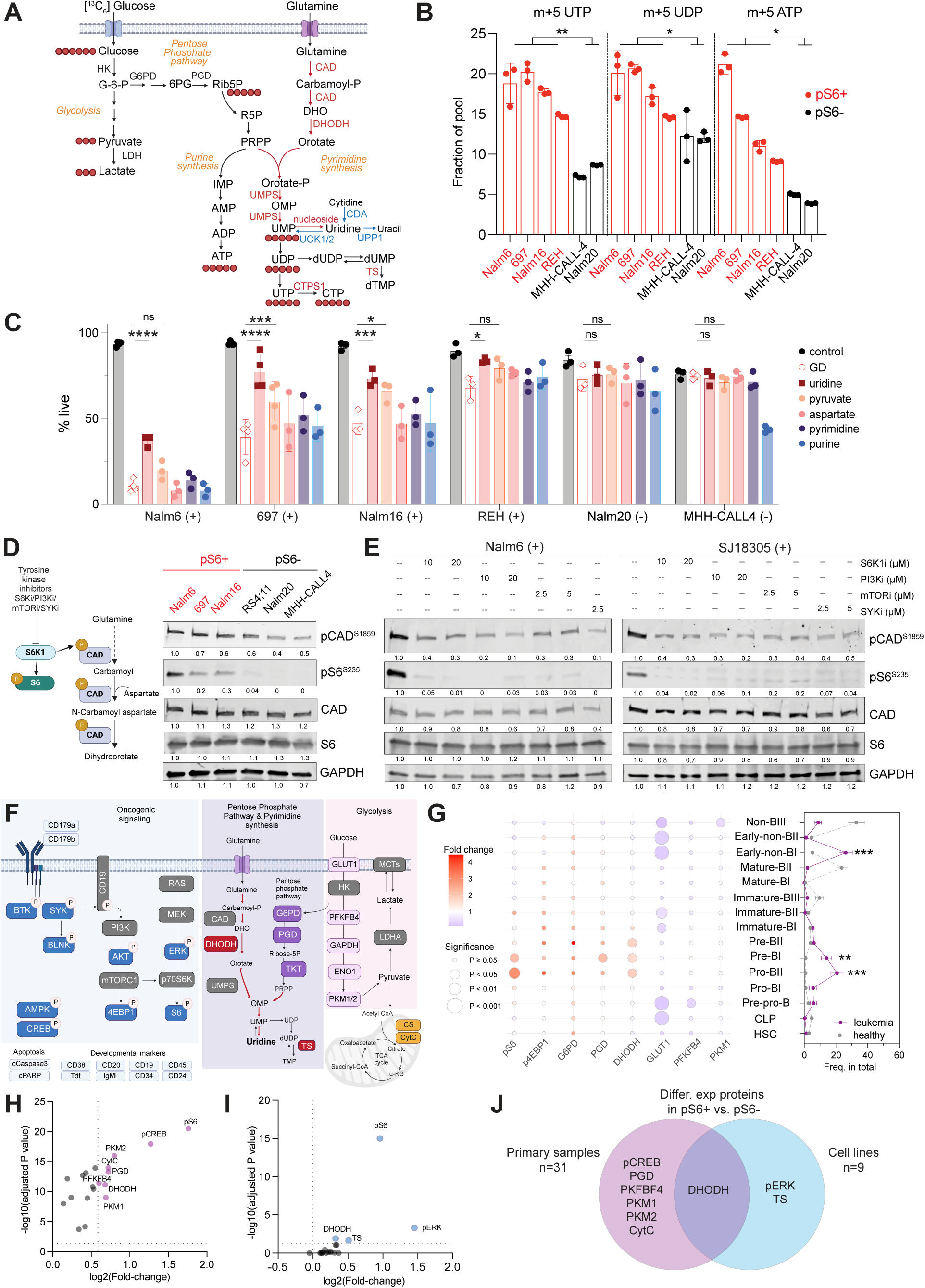
PI3K/mTOR signaling drives glucose-dependent uridine synthesis in B-ALL cells. **A.** Schematic of ¹³C₆ glucose tracing to illustrate glucose flow through glycolysis, pentose phosphate pathway, and purine/pyrimidine synthesis. De novo synthesis converts phosphoribosyl pyrophosphate (PRPP) into uridine monophosphate (UMP) or inosine monophosphate (IMP), while the salvage pathway recycles nucleosides and nucleobases into nucleoside 5′-monophosphates (NMPs) or deoxy NMPs in one adenosine triphosphate (ATP)- or PRPP-dependent step. Pyrimidine synthesis pathways are highlighted: red for de novo, blue for salvage. Key enzymes include CAD (carbamoyl phosphate synthetase II, aspartate transcarbamoylase and dihydroorotase), CTPS1 (CTP synthase 1), DHODH (dihydroorotate dehydrogenase), TS (thymidylate synthase), UCK1/2 (uridine–cytidine kinases 1 and 2), UMPS (UMP synthase); UPP1 (uridine phosphorylase 1). **B.** Fractional labeling of m+5 glucose carbons in pentose phosphate pathway products: UDP (uridine diphosphate, p = 0.0233); UTP (uridine triphosphate, p =0.002) and ATP (adenosine triphosphate, p =0.0342) in glutamine deprived conditions in pS6+ and pS6-cell lines. **C.** Cell viability after glucose deprivation (GD) for 48 hours with and without supplementation of various metabolites: uridine (1mM), pyruvate (5mM), aspartate (5mM), pyrimidine (1mM) and purine (1mM). **D.** Pathway of carbamoyl phosphate synthetase (CAD) phosphorylation (left panel). Western blot of pCAD (S1859), pS6 (S235/S236), total CAD, total ribosomal protein S6 and GAPDH (as loading control) in pS6+ (Nalm6, 697, Nalm16, in red) and pS6-(RS4;11, Nalm20, MHH-CALL4, in black) cell lines. Band intensities were analyzed by Image J software and normalized to first lane and loading control as indicated under each lane. **E**, Western blot of pCAD (S1859), pS6 (S235/S236), total CAD, total ribosomal protein S6 and GAPDH (as loading control) in B-ALL cell line Nalm6 and PDX sample (SJ18305) in the presence or absence of tyrosine kinase inhibitors targeting kinases in the PI3K/mTOR pathway. S6K1 (PF-4708671: 10, 20μM) PI3K (LY294002: 10, 20μM), mTOR (rapamycin: 2.5, 5μM), SYK (PRT062607 HCl: 2.5, 5μM) for 24 hours. **F.** Mass cytometry panel utilized to evaluate expression of glycolysis, PPP, and pyrimidine synthesis proteins along with signaling molecules in primary cells. Measured proteins are in color, non-measured proteins are in gray. **G.** Expression of metabolic proteins between healthy bone marrow (n = 5) cell populations and primary B-ALL bone marrow cells from diagnosis (n = 31). Bubbles colored by fold change of median expression (arsinh transformed) and size indicates P value of the difference. The frequency of each classified subpopulation is indicated to the right. Frequencies summarized as mean ± SEM (B-ALL in purple; healthy control in gray). Pro-BII, P = 0.00024; Pre-BI, P= 0.0036; Early-non-BI, P = 0.000352. **H.** Differential expression of proteins in primary cells gated based on pS6 expression. Proteins in purple are significantly increased in pS6+ cells compared to pS6-cells within Pro-BII and Pre-BI cells from primary patient samples. **I.** Differential expression of proteins in pS6+ (n = 6) compared to pS6-cell lines (n = 3). Proteins in blue are significantly increased in pS6+ cells. **J.** Proteins upregulated in pS6+ cells from primary B-ALL samples (n=31) or B-ALL cell lines (n=9). DHODH is the sole shared protein from both cohorts. Data is displayed as mean ± SD. Statistical test used to compare ^13^C labelling fraction between pS6+ and pS6-cells is Welch t test (B). Statistical test used to compare the rescue effect of metabolites from glucose deprivation (C) and compare pS6+ and pS6-cells (H and I) and compare protein expression in different subpopulations between B-ALL and healthy BM (G, left panel) are two-way ANOVA followed by Šidák’s multiple comparisons test. Statistical test used to compare and to compare the frequency of subpopulations in leukemia samples and healthy bone marrows is Welch’s t test followed by Holm-Šidák method for multiple comparison correction (G, right panel). *p < 0.05, **p < 0.01, ***p < 0.001.

### PI3K/mTOR pathway activation drives uridine synthesis

S6 kinase (S6K1) is a tyrosine kinase situated downstream of the PI3K/mTOR pathway that phosphorylates ribosomal protein S6, thus making pS6 a proxy for PI3K/mTOR pathway activity (Extended Data Fig. 5A). S6 kinase also phosphorylates and activates CAD^19,20^. CAD catalyzes the initial steps in *de novo* pyrimidine synthesis to produce uridine (Fig. 2D). We found higher phosphorylated CAD (pCAD) in pS6+ cells compared to pS6-cells (Fig. 2D). To study if the PI3K/mTOR pathway drives *de novo* uridine synthesis through pCAD, we tested tyrosine kinase inhibitors (TKIs) targeting kinases in the PI3K/mTOR pathway (S6 kinase, PI3K, mTOR, SYK). We confirmed inhibition of downstream signaling nodes using mass cytometry (Extended Data Fig. 5B). We found that CAD phosphorylation is inhibited after treatment with TKIs targeting several levels of the PI3K/mTOR network in pS6+ cell lines and PDXs (Fig. 2E, Extended Data Fig. 5C). To understand how kinase inhibition influences glucose utilization and dependency we evaluated glycolytic activity and glucose sensitivity after TKI treatment *in vitro*. SYK inhibition, one of the upstream proteins in the PI3K/mTOR pathway, significantly decreased ECAR in pS6+ cells, but not in pS6-cells (Extended Data Fig. 5D and 5E). SYK, mTOR, or PI3K inhibition alleviated the glucose dependency in pS6+ cells (Extended Data Fig. 5F). These findings indicate that the active PI3K/mTOR signaling that characterizes pS6+ cells governs glucose dependency driving uridine production by regulation of pCAD.

### DHODH is enriched in primary pS6+ cells and pS6+ cell lines

We evaluated cell phenotype, signaling activity, and metabolic protein expression in 31 primary B-ALL patient samples (Fig. 2F, Supplemental Table 4, 5). Leukemia cells were classified into their developmental state using our developmental classifier^13^ (see methods and gating strategy in Extended Data Fig. 6). We observed enrichment of pro-BII, pre-BI, and early progenitor populations in B-ALL patients compared to healthy BM, (Pro-BII, P = 0.00024; Pre-BI, P= 0.0036; Early-non-BI, P = 0.000352, Fig. 2G, right panel). We observed higher pS6 expression in the pro-BII (FC 2.96, p = 0.0017) and pre-BI populations (FC 2.17, p = 0.0267), but not in early progenitors (FC 1.11, p > 0.999) in leukemic cells compared to healthy BM (Fig. 2G and Extended Data Fig. 7A). Expression of glycolysis pathway proteins (GLUT1, PFKFB4, and PKM1) were similar in healthy and leukemic pro-BII and pre-BI populations (Extended Data Fig. 7B). However, proteins in the PPP (phosphogluconate dehydrogenase, PGD) and pyrimidine synthesis (DHODH), were significantly overexpressed in pro-BII and/or pre-BI B-ALL cells (Fig. 2G and Extended Data Fig. 7B). These results expand our published proteomic signature of pS6+ cells ^13^ by defining their metabolic differences from healthy bone marrow.

Next, we compared the protein expression profiles from pS6+ and pS6-pro-BII/pre-BI cells from primary patient samples. pS6+ pro-BII/pre-BI cells have higher expression of signaling proteins (pCREB), PPP and pyrimidine synthesis pathway proteins (PGD, DHODH), and glycolysis proteins (PFKFB4, PKM1/2) (Fig. 2H). Similar analysis in pS6+ *vs.* pS6-cell lines showed that pERK, DHODH, and thymidylate synthetase (TS) are significantly higher in pS6+ cells (Fig. 2I). DHODH was the only protein common in both sample types (Fig. 2J). DHODH is the rate-limiting enzyme of *de novo* uridine synthesis. These results suggest DHODH may be a promising metabolic target in pS6+ cells.

### B-ALL exhibits reliance on *de novo* uridine synthesis

To understand the potential of targeting pyrimidine metabolism in B-ALL, we evaluated the impact of DHODH knockout in the Cancer Dependency Map (DepMap.org). Compared across 17 types of cancer, B-ALL has the highest dependency on *DHODH* (B-ALL *vs*. all other types, p < 0.001, Fig. 3A) in addition to genes involved in *de novo* pyrimidine synthesis (*UMPS, CAD, TYMS,* and *CTPS1*; Fig. 3B), when compared to solid tumors.

**Fig. 3:**
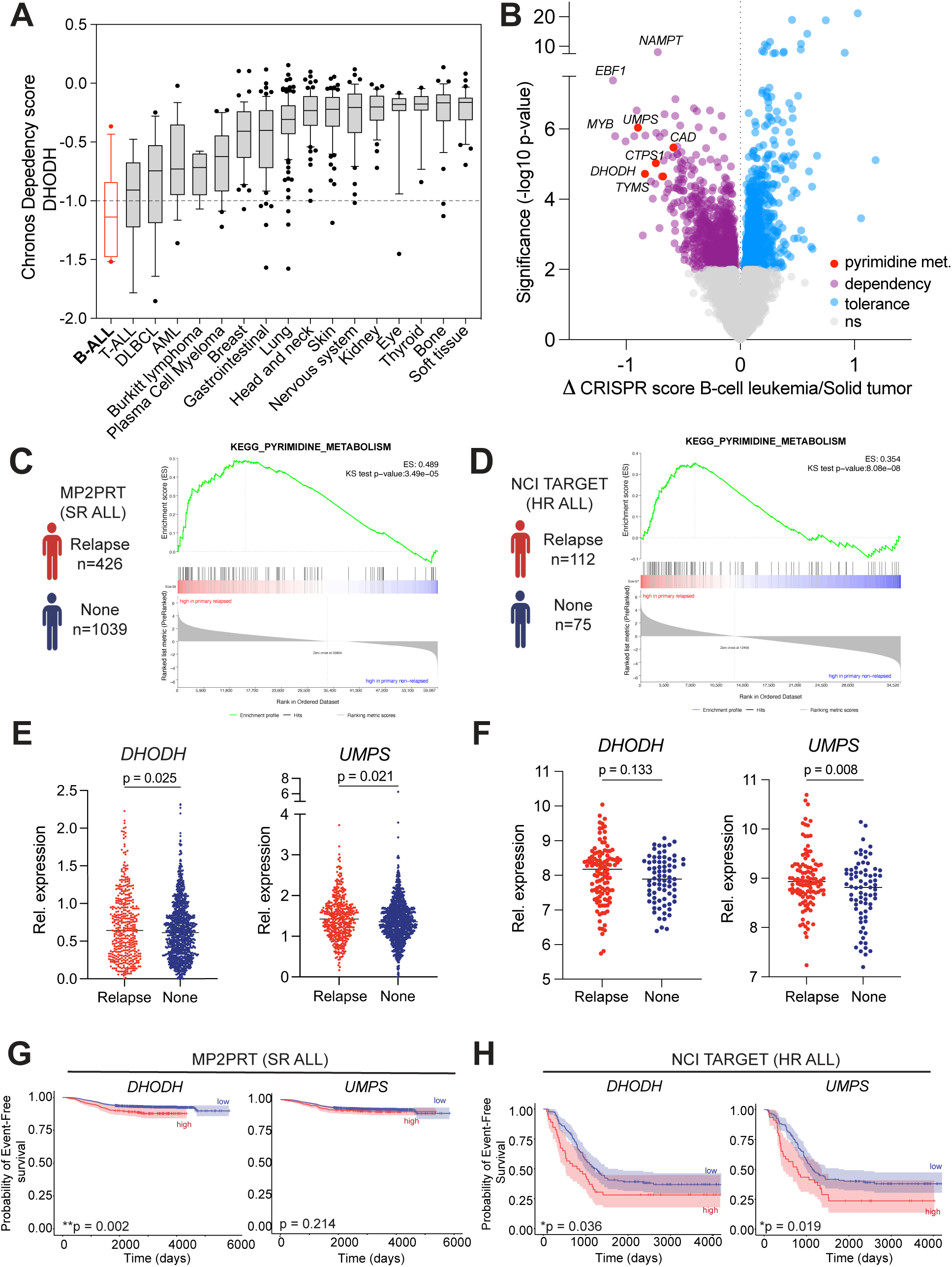
*De novo* uridine synthesis is a metabolic vulnerability in B-ALL. **A.** Effect of DHODH KO in cell lines across cancer subtypes in the genome-wide CRISPR screen (DepMap 22Q2 Public+Score, Chronos). B-ALL (in red) is the most dependent on DHODH for growth and survival among 17 cancer subtypes. The x-axis shows cancer subtypes of cell lines. The y-axis shows the DHODH dependency score (gene effect) per cell line. Commonly essential genes exhibit a median Chronos score of −1 as indicated (dashed line). **B.** Comparison of dependencies in B-cell leukemia (B-ALL) vs solid tumors. The x-axis displays the difference in average CRISPR score per gene between B-ALL and solid tumors, while the y-axis represents significance using -log10(p-value). Purple dots represent “dependency” genes that are preferentially dependent in B-ALL compared to solid tumors. Their loss is more detrimental to B-ALL cells than to solid tumor cells. Blue dots show “tolerance” genes that are not essential in B-ALL. A series of genes involved in pyrimidine synthesis (*DHODH*, *UMPS*, *CAD*, *TYMS* and *CTPS1, red*) are among the most essential genes in B-ALL. **C.** GSEA for KEGG_pyrimidine_metabolism signature between relapsed (n=426) vs non-relapsed (n=1039) patients in the MP2PRT dataset. Enrichment score 0.489, Kolmogorov– Smirnov (KS) test of rank distribution p =3.49e-05. **D.** GSEA for KEGG pyrimidine metabolism signature between relapsed (n=112) vs non-relapsed (n=75) patients with B-ALL in the NCI TARGET dataset. Enrichment score 0.354, KS test of rank distribution p =8.08e-08. **E.** *DHODH* and *UMPS* relative expression in relapsed (n=426) vs non-relapsed (n=1039) patients from MP2PRT dataset (*DHODH*, P=0.025; *UMPS*, p = 0.021). The line indicates mean value. **F.** *DHODH* and *UMPS* relative expression in the diagnostic samples from relapsed (n=112) vs non-relapsed (n=75) patients from NCI TARGET dataset (*DHODH* p =0.133; *UMPS* p = 0.008). The line indicates mean value. **G.** Event-free survival (EFS) based on *DHODH* and *UMPS* expression in MP2PRT dataset when comparing top 10 percentile (N= 147) versus lowest 90% (n=1,318) MP2PRT, Molecular Profiling to Predict Responses to Therapy. Significance determined by Cox regression; *DHODH* p = 0.002; p = 0.214. **H.** EFS based on *DHODH* and *UMPS* expression in the NCI TARGET dataset when comparing the highest quartile (n = 46) to the lowest quartiles (n = 135) Significance determined by Cox regression; DHODH p = 0.036; UMPS p = 0.019. Data in box-whisker plot (A) are shown as median value with the range of values from 10^th^ to 90^th^ percentiles. The dots above or below the lines are outliers. The Kolmogorov–Smirnov test was applied to determine whether the rank distributions of these pathways were statistically different between diagnostic samples from patients who would relapse and patients who are in remission (C and D). Statistical test between relapse vs none was Welch t test. *p < 0.05, **p < 0.01, ***p < 0.001.

Uridine can either be *de novo* synthesized from uridine monophosphate (UMP) or salvaged from cytidine (Extended Data Fig. 8A). In the uridine salvage pathway, cytidine deaminase (CDA) catalyzes the deamination of cytidine to uridine. Uridine is phosphorylated by uridine phosphorylase 1 (UPP1) to fuel PPP and glycolysis. To investigate uridine salvage in B-ALL, we leveraged gene expression data from Depmap. B-ALL cell lines (n=15) had the lowest gene expression of *UPP1* (ranging from 0.09 to 2.4, median 0.29) and *CDA* (ranging from -0.03 to 1.2, median 0.05) among 1,437 cell lines from 31 different cancer subtypes, suggesting that *UPP1* and *CDA* do not regulate uridine utilization and salvage in B-ALL due to lack of expression (Extended Data Fig. 8B and 8C). To determine if B-ALL cells use uridine to generate uracil ribose-1-phosphate to support glycolysis, we attempted to rescue pS6+ cells from glucose deprivation by ribose supplementation. Ribose did not rescue B-ALL cells from cell death in glucose deprived conditions, suggesting uridine is not used to support PPP and glycolysis, in line with the gene expression data (Extended Data Fig. 8D). Alternatively, uridine can be utilized for nucleotide metabolism through its phosphorylation to UMP by the enzyme uridine cytidine kinase 1, encoded by the *UCK1* gene. B-ALL cells display the highest ratio of *UCK1* to *UPP1* among all cancer subtypes, indicating uridine may be preferentially utilized for UMP production (Extended Data Fig. 8E).

### *De novo* uridine synthesis is associated with poor prognosis in B-ALL

Previously, we reported pS6+ cells to be associated with relapse^13^. To investigate the relationship between *de novo* uridine synthesis and relapse in B-ALL, we analyzed two RNA-seq datasets comprising diagnostic samples from children with standard-risk (SR) B-ALL (MP2PRT cohort; n = 1,735)^21^ and high-risk (HR) B-ALL (TARGET cohort; n = 181). Pathway enrichment revealed that diagnostic samples from patients who experienced relapse have higher expression of genes associated with pyrimidine synthesis compared to patients who are in continuous remission (p = 3.49e-05, MP2PRT in Fig. 3C and p = 8.08e-08, TARGET in Fig. 3D). Furthermore, consistent with the gene signatures identified in relapse-predictive pS6+ cells, we observed that expression of genes involved in mitochondrial function, MYC targets, and glycolysis were also higher in the diagnostic samples from patients who experienced relapse (Supplemental Table 6).

Looking at individual genes in the *de novo* uridine pathway, we found that *DHODH* expression was higher in patients who experienced relapse within the MP2PRT dataset (p = 0.025, Fig. 3E). *DHODH* expression predicted inferior EFS in both datasets (MP2PRT p = 0.002, Fig. 3G; TARGET p = 0.036, Fig. 3H) and OS in MP2PRT dataset (p = 0.015, Extended Data Fig. 9A). *UMPS* expression was significantly higher in patients who experienced relapse compared to those in continuous remission within both the MP2PRT (p = 0.021, Fig. 3E) and TARGET (p = 0.008, Fig. 3F) datasets. In addition, higher *UMPS* expression correlated with worse event-free survival (EFS; p = 0.019) and overall survival (OS; p = 0.00026) outcomes among patients in the TARGET dataset (Fig. 3H and Extended Data Fig. 9B). These findings suggest that elevated expression of *de novo* uridine synthesis genes, particularly *DHODH* and *UMPS*, are related to disease prognosis and clinical outcomes.

### Active pS6 signaling predicts sensitivity to DHODH inhibition

To evaluate DHODH activity across different genomic subtypes of B-ALL, we used NetBID2 (data-driven Network-based Bayesian Inference of Drivers)^22,23^ to analyze transcriptomic expression profiles from a published dataset comprising 1,985 B-ALL patients^24^. DHODH was predicted to be active in over half the B-ALL genomic subtypes (54%; Fig. 4A). Encouragingly, we observed high predicted activity of DHODH in several subtypes of high-risk leukemia (e.g. *KMT2A* rearranged, *MEF2D*, *PAX5 P80R*, *BCL2/MYC*)^24,25^. DHODH is also predicted to be active in more common subtypes like *DUX4*, *ETV6-RUNX1,* and *TCF3-PBX1,* while less active in hyperdiploid and hypodiploid subtypes. There was high variability in predicted DHODH activity in Ph-like and Ph+ subtypes, suggesting metabolic heterogeneity in patients within the same genomic subtype and consistent with our single-cell proteomic data.

**Figure 4.**
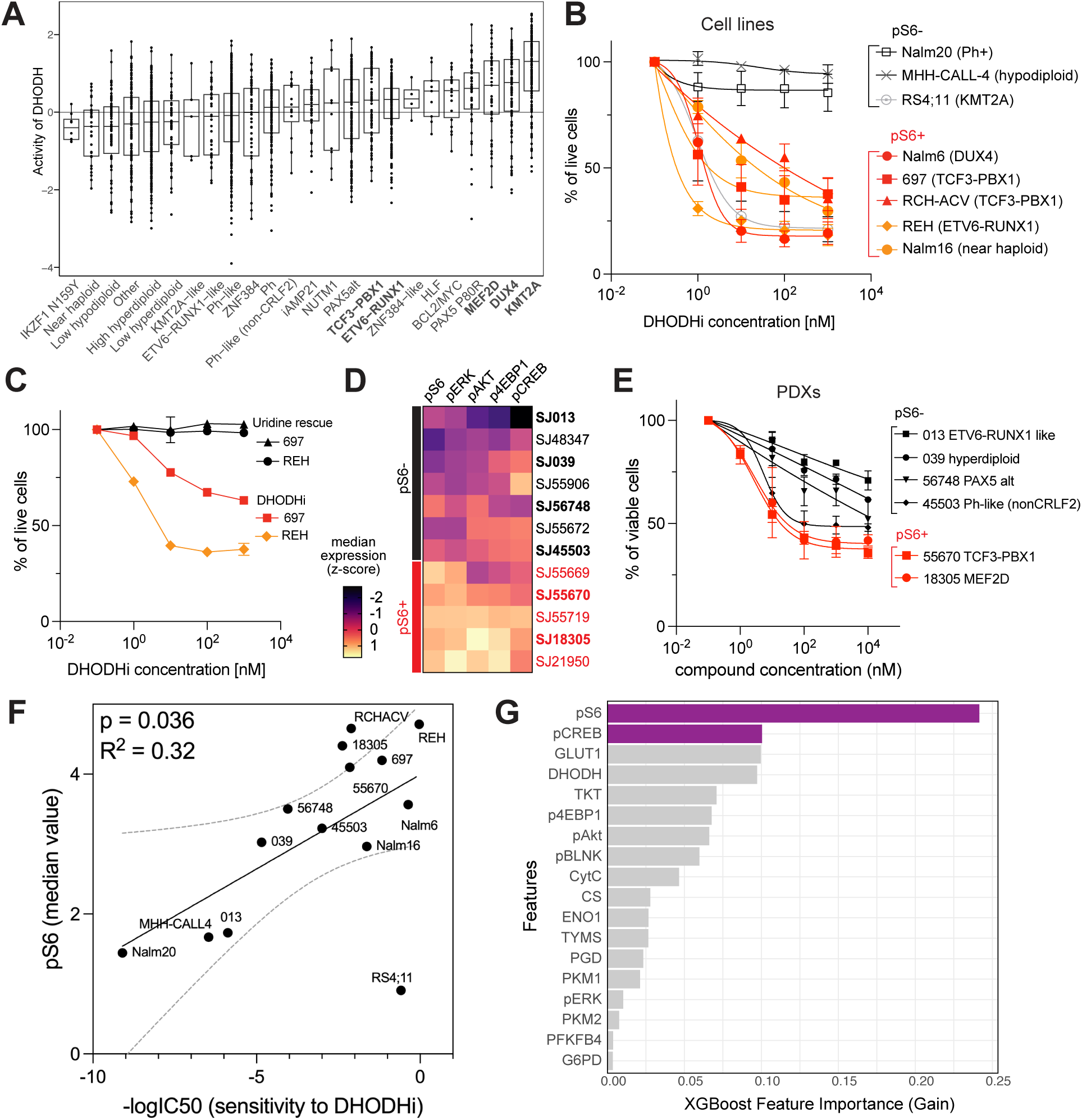
Active pS6 signaling predicts sensitivity to DHODH inhibition. **A.** DHODH activity in 1985 molecularly defined B-ALL cases. Bold indicates subtypes tested for DHODH inhibition (DHODHi) sensitivity in vitro. **B.** In vitro killing after treatment with DHODHi BAY-2402234 in cell lines (n=9). Cells were treated with increasing concentrations of BAY-2402234 for 48 hours. Apoptosis was measured by flow cytometry with Annexin V/7AAD staining. pS6+ magnitude is indicated by color with red/orange representing highest pS6 magnitude (n=5) while pS6-cells (n=3) are indicated in gray. **C.** Viability of REH or 697 pS6+ cells treated with increasing concentration of BAY-2402234 with or without the addition of uridine (1mM) to the cultures over 48 hours. Cell apoptosis was measured by flow cytometry with Annexin V/7AAD staining. **D.** Z-score based on the median expression of signaling molecules pS6, pERK, pAKT, p4EBP1 and pCREB in Pro-BII and Pre-BI cells from patient-derived xenograft (PDX) samples (n=12). Phospho-protein profiles were measured in CyTOF and each cell was classified by developmental classification. We defined cell lines and PDXs with pS6 median expression values (arcsinh transformed) greater than 3 as pS6+ and those no greater than 3 as pS6-. pS6+ PDXs are shown in red, while pS6-PDXs are in black. PDXs marked in bold were used for DHODHi treatment in panel E. **E.** In vitro killing after treatment with BAY-2402234 in PDX samples (n=6). Cells were treated with increasing concentrations of BAY-2402234 for 48 hours. Cell viability was measured by flow cytometry with Annexin V/7AAD. pS6+ PDXs = red; pS6-PDXs = black. **F.** Correlation between the strength of pS6 signaling (median expression) and the sensitivity to DHODHi (-logIC50) in cell lines and PDXs (p = 0.036, R^2^ = 0.32). IC50 values were determined from panel B and E. pS6 median value (arcsinh transformed) was determined by CyTOF. **G.** Cellular features ranked by importance in predicting sensitivity to DHODH inhibition with BAY-2402234 as selected by XGBoost. The top 2 features (> 0.1) are highlighted in the purple. Data in bar graph (A) are shown as median ± SD. Data in curves (B, C, E) are mean ± SD. Linear regression correlation was evaluated in F. The best fit line was shown with 95% confidence bands (dashed curves). *p < 0.05, **p < 0.01, ***p < 0.001.

To validate these predictions and explore the therapeutic potential of DHODH targeting in B-ALL, we assessed the half-maximal inhibitory concentration (IC50) values of DHODH inhibitor BAY-2402234 in B-ALL cell lines of different genomic subtypes (Fig. 4B). Remarkably, DHODH inhibition was effective in nanomolar concentrations. As predicted by NetBID2 analysis, we observed the most pronounced response to DHODH inhibition in cell lines harboring *DUX4*, *TCF3-PBX1*, *ETV6-RUNX1*, and *KMT2A* rearrangements. Notably, Nalm16, which possesses a near-haploid karyotype with TP53 mutation, also demonstrated sensitivity to DHODH inhibition. Furthermore, we observed that pS6+ cell lines were more sensitive to DHODH inhibition, with 5 out of 5 (100%) exhibiting a robust response compared to only 1 out of 3 (33.3%) pS6-cells (Fig. 4B). Uridine supplementation effectively abolished the killing effects of DHODH inhibition, further confirming that uridine is a critical metabolic dependency for pS6+ cells (Fig. 4C).

We then tested a panel of PDX across several genomic subtypes. As in B-ALL cell lines (Fig. 1C), we defined the pS6 signaling strength for each PDX (Fig. 4D). Concordantly with NetBID2 predictions and the cell line results, pS6+ PDXs with MEF2D and TCF3-PBX1 rearrangements had the most pronounced response to DHODH inhibition (Fig. 4E). Notably, we found a robust correlation between the strength of pS6 signaling (median expression) and sensitivity to DHODH inhibition in both cell lines and PDXs (p = 0.036, Fig. 4F). Based on their IC50 values, we categorized cell lines and PDXs as sensitive (IC50 < 1 mM), or resistant (IC50 ≥ 1 mM) to DHODH inhibition. We applied our developmental classification and demonstrated that pro-BII and pre-BI are the most abundant populations in both cell lines and PDXs, in line with patient samples (Extended Data Fig. 10). To identify cellular features predictive of response to DHODH inhibition we used XGBoost (eXtreme Gradient Boosting) in binary classification mode ^26^. We found that pS6 and pCREB expression in pro-BII and pre-BI cells were the most critical predictors of response to DHODH inhibition (Fig. 4G). We confirmed that DHODH inhibition selectively targets pS6+ cells over time at the single-cell level (Extended Data Fig. 11). These data demonstrate pS6 signaling is a surrogate marker for uridine dependency.

### DHODH inhibition prolongs survival in pS6+ B-ALL xenograft models

Given that the strength of pS6 signaling predicts the *in vitro* sensitivity to DHODH inhibition, we assessed the efficacy of *in vivo* treatment with DHODH inhibition in two cell lines and two PDX models with varying pS6 signaling strengths. B-ALL PDX cells (SJ18305, SJ45503) or cell lines (Nalm6, Nalm16) were engrafted in NSG mice and subsequently treated with vehicle or BAY-2402234 (5 mg/kg) five times per week over a total of 24 dosing days (Fig. 5A and Extended Data Fig. 12A). Notably, the 5 mg/kg BAY-2402234 treatment was well-tolerated *in vivo*, with no significant weight loss observed (Extended Data Fig. 12B).

**Figure 5.**
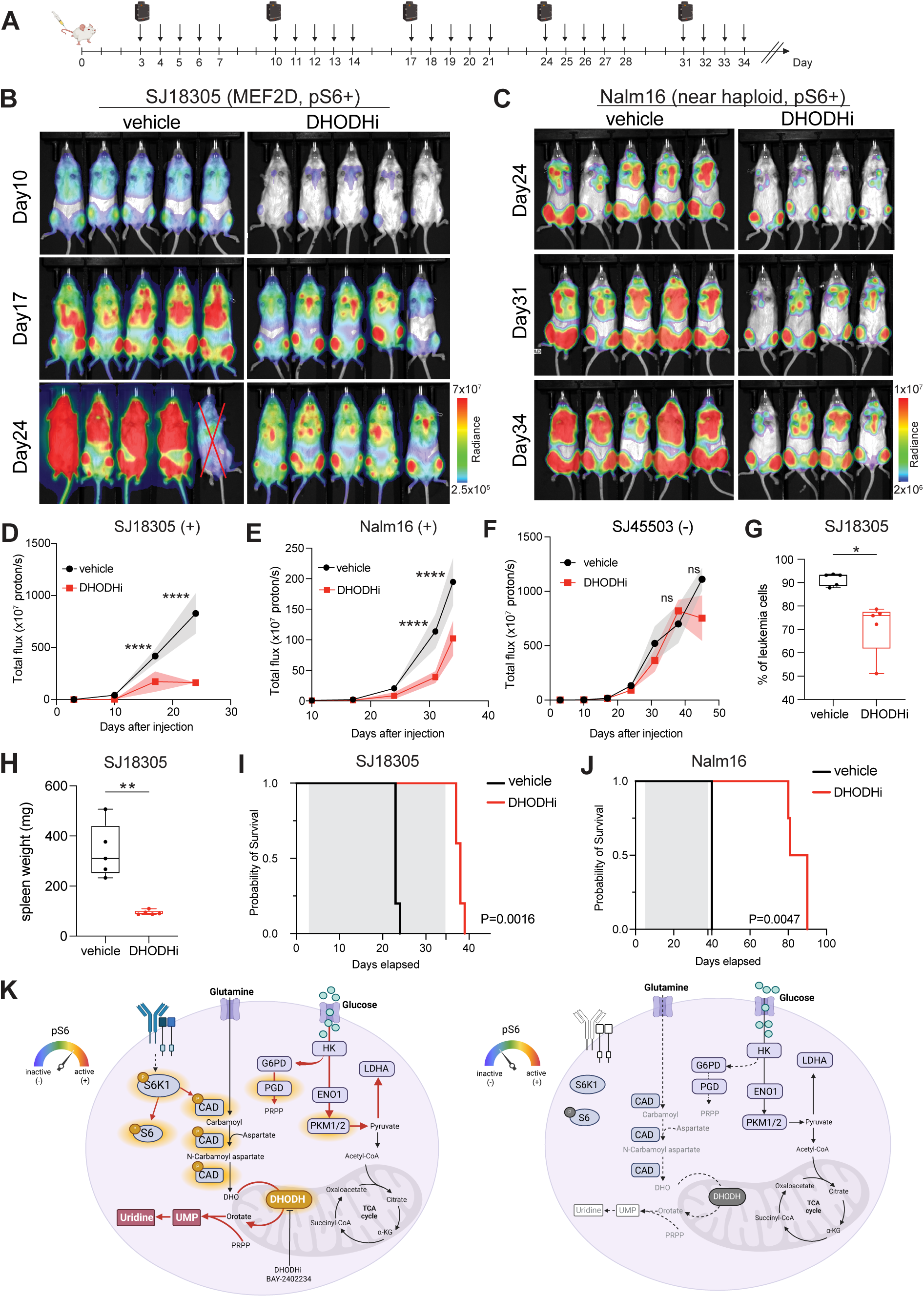
DHODH inhibition prolongs survival of pS6+ B-ALL xenograft models. **A.** Half million B-ALL cells were injected by tail vein in NSG mice. Starting on day +3 following injection, xenografts were treated daily with 5mg/kg BAY-2402234 for 24 dosing days (5 days on, 2 days off). The treatment stopped at 34^th^ day after iv injection. Bioluminescence imaging (BLI) was performed once a week for 5 weeks. **B.** Bioluminescent images of NSG mice at Day 10, Day 17, and Day 24 post engraftment with PDX SJ18305/Luc+ cells. Crossed-out mice indicate experimental mice excluded from the analysis due to death unrelated to leukemia (see methods). **C.** Bioluminescent images of NSG mice at Day 24, Day 31, and Day 34 post engraftment with Nalm16/Luc+ cells. **D.** Leukemia progression in SJ18305 xenografts by bioluminescence in DHODHi (red curve) treated and vehicle (black curve) mice. **E.** Leukemia progression in Nalm16 xenografts by bioluminescence in DHODHi (red curve) treated and vehicle (black curve) mice. **F.** Leukemia progression in SJ45503 xenografts by bioluminescence in DHODHi (red curve) treated and vehicle (black curve) mice. **G.** Leukemia engraftment in DHODHi or vehicle-treated SJ18305 xenografts. **H.** Spleen weight in SJ18305 PDX xenografts in DHODHi treated and vehicle group. **I.** Survival of SJ18305 xenografts treated with DHODHi (red curve) and vehicle (black curve). Gray area indicates the dosing period. **J.** Survival of Nalm16 xenografts treated by DHODHi (red curve) and vehicle mice (black curve). Gray area indicates the dosing period. **K.** Model of uridine dependency and sensitivity to DHODH inhibition in B-ALL. B-ALL cells characterized by active pS6 signaling (left) are glucose dependent for uridine production. Active signaling downstream of PI3K/mTOR pathways activates S6-kinase which phosphorylates CAD, driving uridine synthesis. Consequently, these cells are reliant on de novo pyrimidine/uridine synthesis, making them susceptible to inhibition by targeting DHODH. In contrast, cells lacking pS6 signaling (pS6-) do not depend on uridine synthesis and, therefore, show minimal response to DHODHi treatment. Data in primary samples suggests patients may contain a mixture of pS6+ and pS6-cells but that pS6+, DHODH active cells are associated with chemoresistance. Data in (D) and (E) are mean ± SD tested for significance using a two-way ANOVA mixed model followed by Sidak’s test for multiple comparisons. Data in box plots (F) and (G) are mean ± SD; Welch’s t test was used. Log-rank test was used in Kaplan Meier curves (H). *p < 0.05, **p < 0.01, ***p < 0.001, ****p < 0.0001.

DHODH inhibition significantly slowed tumor progression compared to vehicle in pS6+ samples (SJ18305 p < 0.0001 for both Day 17 and Day 24; Nalm16 p < 0.0001 on Day 31 and Day 34; Nalm6 p < 0.0001 on Day 24 and Day 28; Figs. 5B-E and Extended Data Fig. 12C). By contrast, DHODH inhibition did not slow disease progression in SJ45503, which possessed low pS6 signaling strength (Fig. 5F). In SJ18305, DHODH inhibition significantly reduced leukemia burden within the bone marrow (p = 0.0139, Fig. 5G) and spleen (p = 0.007, Fig. 5H). Survival analysis demonstrated that DHODH inhibition significantly prolonged survival in all pS6+ xenograft models (SJ18305 p = 0.0016; Nalm16 p = 0.0047; Nalm6 p = 0.0031; Fig. 5I-J and Extended Data Fig. 12D) and even demonstrated a modest survival benefit in the SJ45503 mice as compared to vehicle (p = 0.028, Extended Data Fig. 12E). Collectively, these data suggest that targeting uridine synthesis through DHODH inhibition is most effective in ALL cells exhibiting pS6+ signaling yet it has potential therapeutic benefit even in cases with modest pS6 activation.

## Discussion

Relapsed and refractory B-ALL remains a significant clinical challenge, representing the second most common cause of pediatric cancer death. Previous studies have independently associated the activation of B-cell developmental signaling pathways and increased glucose consumption with chemo-resistance and relapse risk^13,27–31^. However, these paradigms have not been previously linked. Here, we demonstrate how activated pS6 signaling leads to a unique metabolic state that promotes glucose utilization for uridine production, uncovering a novel uridine dependency in B-ALL cells. Glucose is converted into uridine through the action of several enzymes in the PPP and *de novo* uridine synthesis pathway, including CAD and DHODH. Gene expression analysis demonstrated the activity of DHODH in most B-ALL genomic subtypes, including high-risk leukemias. Targeting DHODH by pharmacologic inhibition caused cell death *in vitro*, significantly reduced leukemia burden and prolonged survival *in vivo*. Response to DHODH inhibition correlated with the strength of pS6 signaling, suggesting it as a biomarker of uridine dependency in B-ALL. These findings suggest targeting DHODH is a promising therapeutic approach for chemo-resistant B-ALL.

In normal B-cell development, pro-B cell proliferation and survival are primarily driven by IL-7R signaling through the PI3K and JAK-STAT pathways. As development progresses to pre-B cells, IL-7R signaling partners with the pre-BCR, activating RAS and mTOR pathways^14^. Several lines of evidence suggest that sustained signaling of these pathways is associated with chemo-resistance and relapse in B-ALL. Genetic alterations resulting in constitutive signaling in PI3K, mTOR, and RAS pathways characterize high-risk genomic subtypes such as Ph-like and Ph+ ALL^15–17^. Further, activation of SYK, a proximal member of the pre-BCR and PI3K signaling pathways, is thought to contribute to chemo-resistance and relapse in TCF3-PBX1 and KMT2A-rearranged B-ALL^32–34^. We previously described the activation of these signaling pathways as predictive of relapse after chemotherapy^13^. We have also demonstrated that glucocorticoid-resistant B-ALL cells activate these pathways, suggesting that these signaling axes support chemo-resistance^13,30^. However, direct therapeutic targeting of active RAS, PI3K, and mTOR pathways using TKIs has not resulted in significant improvements in disease control, with the exception of BCR-ABL rearranged (Ph+) B-ALL^35^. Multiple mechanisms of resistance to TKIs have been observed, including activation of parallel signaling pathways to bypass inhibition of a single target^36–38^. mTORC1 and S6 kinases are common downstream molecules in several of these pathways and drive *de novo* uridine synthesis via CAD phosphorylation^19,20^. Our results show that pS6+ cells have higher activated phosphorylated CAD and that TKIs targeting various levels of the pathway abolish CAD phosphorylation. CAD catalyzes the initial steps in the *de novo* uridine synthesis pathway, followed by DHODH, which catalyzes the rate-limiting step. Thus, we propose that directly targeting *de novo* uridine synthesis through DHODH inhibition overcomes the challenges of TKI resistance by hitting the convergence of several signaling pathways.

Oncogenic signaling and transcription factor dysregulation in B-cell progenitors induce metabolic changes influencing their transformation^39^. For example, PAX5 and IKAROS exert tumor suppressor functions in B-cell progenitors through transcriptional regulation of glucose transport, glycolysis, and glucose metabolism^40,41^. In addition, increased access to glucose has been associated with chemo-resistance and poor patient outcomes in B-ALL, with inhibition of glycolysis shown to sensitize chemo-resistant cells^28^. Obesity, elevated body mass index or hyperglycemia requiring insulin are associated with higher risk of relapse^42–47^. Further, in line with our data, Xiao et al. determined that B-ALL cells divert glucose-derived carbons toward the PPP^48^. Here, we demonstrate that pS6+ B-ALL cells preferentially utilize glucose for *de novo* uridine synthesis. Primary patient cells with active pS6 signaling have higher expression of DHODH. Further, the expression of DHODH is higher in primary patients who experienced relapse, and DHODH expression alone predicts inferior outcomes in gene expression data from over 1500 patients.

Since Sidney Farber introduced anti-folates in childhood leukemia treatment over 70 years ago^49^, anti-metabolites targeting purine synthesis have been part of therapeutic backbones. However, pyrimidines have not been specifically targeted. While DHODH inhibitors have been approved for use as immunomodulatory agents, their therapeutic potential in cancer is emerging. In solid tumors, DHODH is identified as a target in KRAS-mutant pancreatic adenocarcinoma, IDH-mutant high-grade glioma, MYC-amplified medulloblastoma and neuroblastoma^50–54^. In hematologic malignancies, it has been explored in preclinical settings in AML^55–59^, B-cell lymphoma^60,61^, and T-ALL^62,63^, but clinical trials in AML have not progressed due to lack of efficacy (NCT03404726). Questions remain whether this lack of clinical efficacy was due to suboptimal dosing regimens resulting in a lack of sustained DHODH inhibition or to failure to enrich for patients with subtypes most likely to respond. Our data suggest DHODH is a more promising target for lymphoid malignancies like B-ALL, which exhibit greater dependence on DHODH compared to AML. Analysis of DepMap data highlights B-ALL as the most dependent cancer type on DHODH among 31 cancers. Unlike pancreatic cancer and AML, in which uridine salvage fuels PPP and glycolysis^64,65^, B-ALL demonstrates minimal uridine salvage activity, as indicated by very low expression of UPP1 and CDA. This renders B-ALL more susceptible to DHODH inhibition. Gene expression analysis of 1,985 B-ALL patients showed uridine dependency in most B-ALL genetic subtypes, including high-risk leukemias patients with KMT2A-rearranged, PAX5 P80R, BCL2/MYC^24,25,66^. Consistent with this, DHODH inhibition was detrimental in several *in vitro* and *in vivo* standard-risk and high-risk leukemia models. In cell culture, there is no source for uridine salvage, while *in vivo*, mice and humans can scavenge uridine from dietary sources theoretically inducing resistance against DHODH inhibition over time. Future work will interrogate these mechanisms of resistance and identify agents that enhance response and sensitivity to DHODH inhibition. Given uridine’s critical role in protein glycosylation and phospholipid production, future studies are needed to understand how uridine metabolism contributes to chemo-resistance in B-ALL and how to leverage this metabolic vulnerability for therapeutic advance.

In conclusion, we report a novel vulnerability of chemo-resistant B-ALL cells to uridine. We link activated pS6 signaling in chemo-resistant B-ALL cells with higher activity of *de novo* uridine synthesis. Notably, we found a correlation between the strength of pS6 signaling and sensitivity to DHODH inhibition, highlighting a potential prognostic role for pS6 signaling. This work lays the foundation for future studies to explore DHODH inhibition as a therapeutic approach to relapsed or resistant B-ALL.

## Methods

### Cell culture

697, Nalm6, Nalm16, Nalm20, REH, RS4;11 cell lines were purchased from ATCC (Manassas, VI, USA). RCH-ACV, Kasumi-2 and MHH-CALL-4 were purchased from DSMZ396 (Braunschweig, Germany). 697, Nalm6, Nalm16, REH, RS4;11, RCH-ACV and Kasumi-2 were cultured in RPMI-1640 medium supplemented with 10% fetal bovine serum (FBS). MHH-CALL-4 and Nalm20 were cultured in RPMI-1640 medium supplemented with 20% FBS. For all cell lines, the medium was additionally supplemented with 2mM L-glutamine (Invitrogen) and 1x penicillin/streptomycin (Invitrogen), and cells were maintained at 37 °C and 5% CO_2_.

### Bone marrow samples from patients and healthy donors

Bone marrow samples from healthy donors were purchased from AllCells, Alameda, CA, USA (n = 3; median age was 21 years (range, 19 – 22 years); 2 males and 1 female). De-identified primary patient samples were obtained from the Bass Center Tissue Bank from the Bass Center for Childhood Cancer and Blood Diseases at Lucile Packard Children’s Hospital at Stanford. Thirty-one samples were collected at the time of diagnosis under informed consent. The Stanford University Institutional Review Boards approved the use of these samples. Clinical metadata is available in Supplemental Table 5.

### Patient derived xenograft (PDX) samples

Twelve PDX samples were obtained from the Public Repository of Patient-Derived and Expanded Leukemias (PROPEL; propel.stjude.cloud). Details regarding these PDX samples are available in Supplemental Table 7. Six of twelve PDX samples were able to be cultured without significant cell death in vitro in StemSpan™ SFEM (StemCell technologies) for a week.

### Definition of pS6+ and pS6-cell lines and PDXs

The pS6 signaling strength was defined as the arcsinh-transformed median expression value of pS6 profiled by CyTOF (see Supplemental Methods). As a cutoff, we defined cell lines and PDXs with pS6 median values greater than 3 as pS6+ and the ones no greater than 3 as pS6-.

### Extracellular flux analysis

To profile the glycolytic and mitochondrial activity in cell lines, the oxygen consumption rate (OCR) and extracellular acidification rate (ECAR) were measured using a Seahorse XFe24 analyzer (Agilent). Briefly, a sensor cartridge (102342-100, Agilent) was hydrated in a Seahorse XF Calibrant (100840-000, Agilent) overnight at 37°C in a non-CO_2_ incubator. On the day of measurement, cells were collected and washed with PBS. After centrifugation, cells were resuspended at 2×10^6^ per mL in Seahorse XF RPMI medium (pH 7.4) supplemented with 10mM glucose, 2mM glutamine and 1mM pyruvate. 100 μl cell suspension was added into each well of Seahorse XF Cell Culture Microplate (102342-100, Agilent). After centrifugation at 200 × *g* for 2 min with no brake, the plate was equilibrated for 30 min in a 37°C incubator without CO_2_. Additional 500ul medium were added and incubated for 30 min before loading to the machine. The OCR and ECAR were measured in basal conditions.

### Nutrient deprivation, metabolite rescue and cell viability assay

To test the nutrient dependencies of glucose and glutamine in cell growth and survival, cell lines were cultured under regular conditions, glucose or glutamine deprivation conditions at 37 °C, 5% CO_2_ for 48 hours. Cells were stained with Annexin V (Pacific Blue, Biolegend, 640918) and 7AAD viability staining solution (Biolegend, 420404) for 15 min following manufacturer instructions and directly acquired on CytoFLEX Flow Cytometer (Beckman Coulter). Cells negative for Annexin V and 7-AAD are defined as viable cells. Biological triplicates of experiments were performed.

To rescue the effect of glucose deprivation (GD), cells were seeded in 96-well plates at 3-5 × 10^4^ cells in 200 μl of growth medium and supplemented with uridine (1mM), pyruvate (5mM), aspartate (5mM), pyrimidine (1mM) and purine (1mM) under glucose deprivation condition (10% dialyzed FBS, 0mM glucose, 2mM glutamine) every 24 hours at 37 °C, 5% CO_2_. After 48 hours, cell viability was measured by Annexin V/7AAD staining in FACS as above. The chemical supplements were purchased from Millipore Sigma with details shown in Supplemental Table 8.

To evaluate the *in vitro* killing effect of the DHODH inhibitor, cell lines and PDX cells were seeded in flat bottom 96-well plates at 3-5 × 10^4^ per well in 200 μl of growth medium and treated with five increasing doses of DHODHi (BAY-2402234) on a logarithmic scale at 37 °C, 5% CO_2_. After 48 hours, 100ul of cell suspension was transferred to a U-bottom 96-well plate followed by centrifugation and PBS supplemented with 5% FBS wash twice. Cell viability was measured by Annexin V/7AAD staining in FACS as above.

### Stable Isotope Tracing and Metabolic Analyses

B-ALL cell lines (Nalm6, 697, REH, Nalm16, Nalm20 and CALL4 were pretreated with or without glutamine for 4 hours. Following the procedures as previously published^67^, the culture medium was replaced with RPMI1640 with or without L-glutamine (Corning, 25-005-CI) and supplemented with 10 mmol/L U-^13^C_6_-glucose (Cambridge Isotope Laboratories, CLM-1396) and 10% dialyzed FBS (Gibco, 26400044). The cells were incubated for 4 hours under these conditions. After incubation, the cells were washed twice with warm PBS, followed by the addition of ice-cold 80% methanol. The cells were then vortexed briefly and incubated on ice for 15 min. The solution was centrifuged at 15,000g for 15 min at 4°C. The resulting supernatant was collected for LC-MS analysis. Metabolomics and isotope tracing analyses were performed using an Agilent 1290 Infinity Liquid Chromatography (LC) System coupled to a Q-TOF 6545 mass spectrometer (MS; Agilent). Targeted analysis, isotopologue extraction, and natural isotope abundance correction were conducted using Agilent Profinder B.10.00 software as previously described^67^. Data are presented as mean ± SD across three biological replicates.

### Signaling inhibition using tyrosine kinase inhibitors

Cells were incubated with RPMI1640 medium supplemented with 10mM glucose, 2mM glutamine and without FBS overnight at 37°C and 5% CO_2_. After 16 hours of serum starvation, cells were seeded at the density of 1 × 10^6^ per mL in 10mL of growth medium and treated with different concentrations of tyrosine kinase inhibitors to target S6K1 (PF-4708671: 10, 20μM) PI3K (LY294002: 10, 20μM), mTOR (rapamycin: 2.5, 5μM), SYK (PRT062607 HCl: 2.5, 5μM). After 24hrs, cells were washed in PBS and pelleted into two different Eppendorf tubes. One million cells were analyzed by CyTOF and the remaining cells were used to perform western blot.

### Manual gating

Single cells were gated using Omiq software (https://www.omiq.ai/) based on event length and 191Ir/193Ir or 103Rh DNA content to filter out debris and doublets, as previously described^68^. After gating for single cells, live non-apoptotic cells were identified by gating on cleaved poly(ADP-ribose) polymerase (cPARP), cleaved caspase-3 (c-Caspase3), and 195Pt levels^69^. In PDX samples, murine cells were excluded by gating for mouse CD45 (mCD45). Platelets and erythrocytes were removed by gating on CD61 and CD235a, while T cells and myeloid cells were excluded based on CD3e, CD33, and CD16 expression. CD38^high^ plasma cells were also gated out, leaving a population defined as lineage-negative blasts (Lin− B+). Unless noted otherwise, further analysis was performed on this Lin− B+ population.

### B cell developmental classification

We utilized the single-cell developmental classifier previously reported^13^. In summary, the Lin−/ B+ fraction from healthy human BMs was manually gated into 15 developmental populations of normal B lymphopoiesis, mixed progenitors, mature B and non-B cell fractions, as depicted in Supplemental Fig 6. The classification of each group was determined by the expression of 10 key B-cell developmental markers: CD19, CD20, CD24, CD34, CD38, CD45, CD179a, CD179b, intracellular IgM (IgMi), and terminal deoxynucleotidyl transferase (TdT). Using the tidytof R package^70^, we first generated healthy-fit objects by calculating the mean and covariance matrix for all healthy populations. Lin−/ B+ cells from primary leukemia or PDX samples, or live B-ALL cells from cell lines were assigned to the most similar healthy developmental subpopulations based on the shortest Mahalanobis distance across all 10 dimensions. Cells were labeled as ‘unclassified’ if no Mahalanobis distances fell below the threshold (distance = 10, determined by the number of dimensions).

### Depmap genome-wide CRISPR screen data analysis

The CRISPR 23Q4 public data from the screens published by Broad Achilles and Sanger Score projects^71^ were downloaded from the Depmap portal https://depmap.org/portal/download. The CRISPR 23Q4 screening was performed for 31 tumor lineages on 1091 cancer cell lines, of which 12 were annotated as B-ALL. We defined cell lines as solid tumor if they were not annotated as lymphoid or myeloid or non-cancerous. 975 cell lines were assigned as solid tumors. The gene effect scores summarizing the guide depletion were determined based on the Chronos algorithm^72^. The Chronos dependency score lower than -0.3 indicates inhibition of cell growth or death after gene knockout (KO). Commonly essential genes exhibit a median Chronos score of −1. To determine genes that are differentially dependent between B-ALL cell lines and solid tumor lines, Chronos scores were compared, and Welch’s t-test was conducted for each gene between the two groups. Genes are considered significant if their p-value is less than 0.01 (i.e., -log(p-value) > 2). A mean CRISPR score difference below zero indicates that a gene is specifically dependent on B-ALL, while a mean difference of zero or higher identifies the gene as specifically dependent on solid tumors.

## Data analysis in published datasets

### Molecular Profiling to Predict Responses to Therapy (MP2PRT)

We leveraged RNA-seq data from 1,465 diagnostic samples from patients with predominantly SR B-ALL enrolled across four Children’s Oncology Group (COG) clinical trials as published by Ti-Cheng et al^21^. Gene-level summed TPM serve as the metric for GSEA analysis using fsea (version 1.28.0) R package. The Kolmogorov–Smirnov test was applied to determine whether the rank distributions of these pathways were statistically different between the two groups. Log2TPM was used to compare the expression of *DHODH* and *UMPS* between relapse (n=426) and non-relapse samples (n=939) using Welch’s t-test.

For survival analysis, we categorized the patient samples based *DHODH* and *UMPS* expression levels. The high-expression group comprised 147 samples in the highest 10th percentile, while the low-expression group included 1318 samples in the lowest 90th percentile. Since relapse samples were enriched in the datasets, to account for the sample selection probabilities, we adjusted the Kaplan-Meier curves with inverse probability weights. Specifically, the weights are 2.12 for relapse, MRD negative; 2.26 for relapse, MRD positive; 13.86 for non-relapse, MRD negative; and 4.99 for non-relapse, MRD positive. In addition, we used weighted Cox-regression tests to derive the P values.

### Therapeutically Applicable Research to Generate Effective Treatments (TARGET)

We focused on the primary patient cohort (n = 187) and performed gene set enrichment analysis (GSEA) to evaluate whether pre-defined gene sets associated with various metabolic pathways— such as signaling, glycolysis, and pyrimidine synthesis—differed significantly between relapse and non-relapse samples, using the fgsea (version 1.28.0) R package. The Kolmogorov–Smirnov test was applied to determine whether the rank distributions of these pathways were statistically different between the two groups.

In addition, we specifically compared the normalized gene expression levels of *DHODH* and *UMPS* between relapse and non-relapse samples using Welch’s t-test, providing a focused analysis of genes involved in pyrimidine metabolism.

For survival analysis, we categorized the patient samples based on the expression levels of *DHODH* and *UMPS*. The high-expression group comprised 46 samples in the highest 25th percentile, while the low-expression group included 135 samples in the lowest 75th percentile. Kaplan-Meier curves were plotted to compare the probability of event-free survival between the high- and low-expression groups. To analyze the survival curves, we employed the Cox regression test, allowing us to assess the impact of gene expression on patient outcomes.

### NetBID2 analysis to query DHODH activity in patients with B-ALL

We utilized the network-based integrative NetBID2 algorithm^22,23^ to infer DHODH gene activities using bulk RNA-seq data from a published RNA-seq dataset comprising 1,985 B-ALL patients^24^. Briefly, we employed the SJARACNe algorithm to reverse-engineer a B-ALL interactome (BALLi) from this published RNA-seq dataset^24^. To ensure robust network quality, we excluded subtypes with fewer than 15 samples. This analysis targeted 10,843 hub genes, including 1,937 transcription factors and 8,906 signaling proteins. The parameters for SJARACNe were configured as follows: 1) Bootstrap p-value threshold: p = 1 × 10^-7; 2) Consensus p-value threshold: p = 1 × 10^-5; 3) Data Processing Inequality (DPI) tolerance: e = 0; 4) Number of bootstraps (NB): 100. The resulting data-driven BALLi consisted of 33,237 nodes and 314,914 edges. Subsequently, we used the cal.Activity function with parameters es.method = “weightedmean” and std = TRUE in NetBID2 to infer the activity of all hub genes, including DHODH, in each B-ALL patient based on their gene expression profiles. The NetBID2 package is available online at: https://github.com/jyyulab/NetBID.

### Single cell feature selection

Following DHODHi treatment, IC50 values at 48-hour were calculated in GraphPad Prism 10 using nonlinear regression model with “log(inhibitor) vs response -- variable slope (four parameters)” function. Cell lines with logIC50 values below 3 (IC50 < 1 μM) are categorized as sensitive (response to DHODHi), whereas cell lines with logIC50 values equal to or more than 3 (IC50 ≥ 1 μM) are grouped as resistant lines. Protein profiles from cell lines and PDX samples were acquired by CyTOF. We performed batch effect correction and expression normalization using the limma R package. After developmental classification of cell line and PDX samples, we focused on the most dominant subpopulations: pro-BII (n= 139,646) and pre-BI (n= 71,675). Cells were grouped into sensitive (n=147,245) or resistant (n=64,076) groups and concatenated respectively. We split this data into training (80%) and test set (20%) and developed XGBoost (eXtreme Gradient Boosting) binary classification model^26^ to retrieve the importance score for each feature as profiled by CyTOF.

### GFP Expression in PDX cells

293GP were used (generously gifted by the laboratory of Dr. Garry Nolan) for retrovirus production. Briefly, 293GP cells at 70% confluency on 10 cm Poly-D-Lysine coated plates were co-transfected with 9 μg of luciferase-mNeoGreen vector and 4.5 μg RD114 envelop plasmid (graciously provided by Dr. Crystal Mackall) with TurboFect (Thermo Fisher Scientific). Viral supernatants were collected at 48 and 72 hours post transfection by centrifuging to pellet cell debris and stored at -80°C. Nalm6, Nalm16 and PDX samples (SJ18305, SJ45503) cells were transduced with retroviral supernatant. Briefly, 5 μg/ml vitronectin in PBS (Takara) was used to coat non-tissue culture treated 12-well plate at 4°C overnight. The next day, wells were washed with PBS and blocked with PBS supplemented with 2% BSA for 15 min. 1 ml of thawed retroviral supernatant was added to the well and centrifuged at 3,200 rpm for 1.5 hours, followed by adding 5 × 10^5^ cells in each well. 48 hours later, transduction efficiency was evaluated by GFP expression in CytoFLEX instrument (Beckman Coulter). Transduced cells were enriched by sorting GFP-positive cells in a BD FACSAria II Cell Sorter (BD Biosciences). Following sorting, cell lines were cultured in 10% FBS RPMI1640 medium and PDX cells were cultured in StemSpan™ SFEM media as isogenic cells and tested for *Mycoplasma* contamination for future use *in vivo* studies.

### *In vivo* DHODHi treatment in cell line and PDX models

NOD/SCID/IL2Rγ^−/-^ (NSG) mice were purchased from the Jackson Laboratory, housed, and treated under the Stanford University Committee on Animal Welfare-approved protocol. Six-to-eight week old female mice were engrafted with 5.0 × 10^5^ cells (Nalm6-Luc^+^, Nalm16-Luc^+^, SJ18305-Luc^+^ or SJ45503-Luc^+^) via intravenous (I.V.) injection. When engraftment was detectable by bioluminescence imaging (BLI) (BLI > 1 × 10^6^), mice were randomized in two different experimental groups: vehicle (5% DMSO, 40% PEG400, 5% Tween 80, 50% saline) and DHODHi BAY2402234 (MedChemExpress). Two different *in vivo* experiments were performed for PDX xenografted mice and cell line xenografted mice. For PDX xenografted mice, n=5 mice were in the vehicle group; n=10 mice were in the DHODHi group. Among them, 5 mice were used to perform the survival analysis and the remaining 5 mice were used to compare the leukemia progression and sacrificed simultaneously as mice in the vehicle group. For mice xenografted with cell lines, 5 mice were used for each group to assess engraftment and survival. From 3- or 5-days until 34- or 38-day post engraftment, mice received 5 mg/kg (5 days per week) of DHODHi or vehicle via oral gavage (O.G.). Engraftment was monitored once or twice per week by bioluminescence (BLI) analysis and was assessed as the percentage of hCD19/hCD45+/mCD45.1-cells in peripheral blood. Mice were sacrificed when clinical signs of leukemia were observed. In the survival analysis, mice were censored if they were moribund for other reasons, such as accidental death (n = 1 in the Nalm6 vehicle) or did not develop leukemia (n = 1 in the Nalm16 treatment group). Whensoever suspected leukemia-unrelated deaths occurred, FACS analysis of bone marrow and spleen was performed to confirm that death was not related to a high burden of leukemia.

### Statistical analysis

Data were analyzed and visualized using R statistical software (http://www.r-project.org) or GraphPad Prism 10 software. P values were calculated using the statistical test described in the relevant figure legend. P < 0.05 was considered statistically significant, and P values are denoted with asterisks as follows (P > 0.05, not significant, n.s.; *, P < 0.05; **, P < 0.01; ***, P < 0.001; and ****, P < 0.0001).

## Data availability

Mass cytometry data from clinically annotated patient samples, cell lines, and PDX samples are available in Community Cytobank (https://community.cytobank.org).

Bulk RNA-seq data from primary samples in our previously published patient cohort (n=6) have been deposited in NCBI Gene Expression Omnibus (GEO) and are accessible through GEO.

The TARGET dataset used for this study is accessible through the TARGET website at https://ocg.cancer.gov/programs/target/data-matrix. TARGET BAM and FASTQ sequence files are accessible through the database of genotypes and phenotypes (dbGaP; https://www.ncbi.nlm.nih.gov/projects/gap/cgi-bin/study.cgi?study_id=phs000218.v24.p8) under accession no. <u>phs000218</u> (TARGET) and at NCI’s Genomic Data Commons (http://gdc.cancer.gov) under project TARGET. Transcriptomic data in MP2PRT dataset are accessible in dbGaP: Project ID: MP2PRT-ALL; accession number: phs002005.v1.p1 (https://www.ncbi.nlm.nih.gov/projects/gap/cgi-bin/dataset.cgi?study_id=phs002005.v1.p1).

## Acknowledgments

We would like to thank current and past members of the Davis laboratory for helpful discussions. This work was supported by the National Institutes of Health R01-CA251858 and the Stanford Maternal Child Health Research Institute, MCHRI. Y.L. is supported by R01-CA251858 and MCHRI. J.S. is supported by Associazione Italiana per la Ricerca sul Cancro (AIRC; grant no. 27325). K.L.D is supported by the Anne T. and Robert M. Bass Endowed Faculty Scholar in Pediatric Cancer and Blood Disease and the Harriet and Mary Zelencik Endowment. We acknowledge the PROPEL project at St. Jude for contribution of annotated PDX expanded ALL cells.

## Author contributions

Y.L. conceived of, designed, and led this study, designed and performed experiments, analyzed and interpreted data, generated the figures, and wrote the manuscript. H.J. conducted experiments and analyzed stable isotope metabolomics data. J.L. performed data analysis for the TARGET and MP2PRT datasets. L.S., A.J., and D.J. contributed to mass cytometry experiments and validated the antibody panel used in these experiments. M.M assisted with *in vivo* mouse studies, cell culture work, retrovirus generation and transduction. A.K. performed the CyTOF data analysis using XGboost. T.C. provided guidance for MP2PRT dataset analysis. P.D. and J.S. performed cell sorting, RNA extraction and performed RNA-seq data analysis. J.M assisted in genomic analyses. T.K. contributed to developmental classification data analysis using tidytof. A.W. contributed to western blot analysis. F.H. and S.B. provided guidance on mass cytometry experiments. M.H., N.J.L, K.M.S., C.G.M., M.L., J.Y. and J.Y. contributed patient or PDX samples and provided clinical data. H.J., J.L., L.S., P.D., J.Y., J.Y. and J.Y. provided guidance and scientific input. K.L.D. conceived of and designed this study, provided guidance and scientific input, interpreted data, and wrote the manuscript. All authors discussed the results and prepared the manuscript.

