## Supplemental methods and figures for "Uridine Metabolism as a Targetable Metabolic Achilles’ Heel for chemo-resistant B-ALL"

**Cell sorting to enrich pre-B cells from patient samples**

Six diagnostic samples of known pS6 status from our previously published cohort[1] (n=3 pS6+ from patients who would relapse; n=3 pS6- from patients who are in continuous remission) were thawed in warm RPMI-1640 medium with 10% FBS. The details about patients were listed in **Supplemental Table 1**. To enrich leukemic pre-B cells from patient samples, samples were stained with 7-color FACS panel listed in **Supplemental Table 2** for 30min at room temperature. Pre-B cells are enriched in the population with low CD45 expression, positive for CD34, CD38 and CD10 and high expression of CD19, CD24 and CD43. Sorted cells were collected in 1 mL of cell-staining media containing RNAase inhibitors (VRC, Promega; RNAsin, New England Biolabs) and RNA was extracted using RNeasy Micro kit (Qiagen) following manufacturer's instructions.

**Bulk transcriptomic sequencing analysis**

**Enriched pre-B cells:** RNA samples were submitted for library preparation and sequencing to SFGF (Stanford Functional Genomics Facility). cDNA libraries were generated using the SMARTer Ultra Low kit (Takara Bio/Clonotech Laboratories) and a low-input library kit was used for library generation before being sequenced using Illumina HiSeq 4000 system. After quality check, FASTQ files were first pre-processed using fastp to perform quality profiling, read filtering, and adapter trimming. Afterward, clean reads were aligned to the human reference genome (hg38.v93) and salmon was used to quantify transcript abundance[2]. Then, the transcript level expression data were summarized to the gene level using tximport R package[3] low-expressed genes (less than 10) were removed before proceeding with data analysis. Normalized counts were analyzed by using DESeq2 package in R[4]. Gene Set Enrichment Analysis (GSEA) was performed by GSEA software v4.1.0 (BROAD Institute—UC San Diego joint project) using default settings. The Hallmark database was used for the analysis and the normalized enrichment score (NES) was plotted by using GraphPad Prism.

**MP2PRT dataset analysis**: Briefly, FASTQ files of these samples were pre-processed for quality profiling and read filtering. The adapters in sequencing reads were trimmed with Trim Galore (v0.4.4). The trimmed sequencing reads were mapped with STAR[5] to human genome GRCh38. TPM (transcripts per kilobase million) counts were generated using RSEM package.

**TARGET dataset analysis**: The results published here are in part based upon data generated by the TARGET (https://www.cancer.gov/ccg/research/genome-sequencing/target) initiative, phs000463 and phs000464[6]. The data used for this analysis are available at the Genomic Data Commons (https://portal.gdc.cancer.gov) and are approved for this use. In total, there were 187 B-ALL patients included in the dataset. Of these, 46 patients had both diagnosis and relapse samples available, 66 patients experienced relapse, and 75 patients did not experience relapse. Fastq or BAM files from the 187 diagnostic samples and 46 paired relapse samples were downloaded and pre-processed using fastp for quality profiling, read filtering, and adapter trimming. After pre-processing, the clean reads were aligned to the human reference genome (hg38), and salmon was used to quantify transcript abundance. Next, the transcript-level expression data were aggregated to the gene level using the tximport R package. Genes with low expression (fewer than 10 counts) were filtered out. Principal component analysis (PCA) identified batch effects in the samples that were sequenced on different sequencing machines. To address this, we performed batch effect correction and gene count normalization using the limma R package. The normalized gene counts were then used for subsequent analyses.

**Mass cytometry (CyTOF)**

We processed the samples as per the previously published methods[7]. To thaw viable healthy BM, leukemic primary, and PDX cells, we used 90% RPMI medium (Thermo Fisher Scientific) with 10% FCS, along with 20 U/ml sodium heparin (Sigma-Aldrich), 0.025 U/ml Benzonase (Sigma-Aldrich), 2mM L-glutamine, and 1x penicillin-streptomycin (Invitrogen). The cells were rested at 37°C for 30 minutes. For B-ALL cell lines, the cells were suspended in the corresponding culture media. Meanwhile, 1 x $10^6$ healthy BM, leukemic primary, and PDX cells were stained for viability using cisplatin as described in previous work [8]. Following viability staining, cells were fixed with 1.6% paraformaldehyde (PFA, Electron Microscopy Sciences) for 10 min at room temperature. Barcoding was done using in-house prepared 20-plex palladium barcoding plates[9], with one healthy BM sample per plate to control for batch effects. A total of 8 barcode plates were used. Following barcoding, cells were pelleted and washed with cell-staining medium (CSM; PBS with 0.5% BSA and 0.02% sodium azide) to remove residual PFA. Blocking was performed with Human TruStain FcX (BioLegend), following the manufacturer's instructions. Antibodies to surface markers were added to 800 µl final reaction volumes, and samples were incubated at room temperature for 30 min at 300 rpm (**Supplemental Table 4**). Cells were washed with CSM and permeabilized with methanol at 4°C for 10 min. After permeabilization, cells were washed with CSM and stained with intracellular-marker and phospho-specific antibodies in 800 µl for 30 min at 300 rpm (**Supplemental Table 4**). Cells were washed once in CSM, then stained overnight with 1:5,000 191Ir/193Ir or 103Rh DNA intercalator (Standard Biotools) in PBS supplemented with 1.6% PFA at 4ºC. Before acquisition, cells were washed once with CSM, washed twice with double distilled water, filtered to remove aggregates, and resuspended in water with 139La/142Pr/159Tb/169Tm/175Lu normalization beads[10]. The samples were kept on ice and introduced into a Helios mass cytometer (Standard Biotools) at a constant rate of 150 - 200 cells/s using a super sampler.

**Processing of mass cytometry data**

Data were normalized together using bead normalization[10], and files were debarcoded as described[9]. Single-cell protein expression data were extracted and analyzed using packages from the Comprehensive R Archive Network (CRAN) project (https://cran.r-project.org/) and Bioconductor (http://www.bioconductor.org). Raw data were transformed using the hyperbolic arcsine (arcsinh) function with a cofactor of 5. The percentage of positive cells for each phosphorylated protein was calculated based on a mass cytometry cutoff of ≥10 counts. The expression of proteins in each population of interest was determined by calculating the mean or median expression level after arcsinh transformation.

**Cytoplasmic protein extraction and western blotting**

Cytoplasmic fraction of proteins was extracted by NE-PER™ Nuclear and Cytoplasmic Extraction Reagents (Thermo Fisher Scientific) containing protease and phosphatase inhibitors following manufacturer instructions. The concentration of proteins was measured by Pierce™ BCA Protein Assay Kit (Thermo Fisher Scientific). A total of 30 µg of protein was denatured in Laemmli buffer at 95°C for 5 min and Western blotting was performed using the Bio-Rad system (TGX 4–15% gels). Transfer was performed using the Trans Blot turbo system (Bio-Rad) onto PVDF membranes. Primary antibodies for phospho-CAD (S1859), total CAD, total ribosome protein S6, phospho-S6 (S235/236) and GAPDH were purchased from Cell Signaling Technologies. Secondary antibodies (IRDye® 800CW goat anti-rabbit and anti-mouse IgG) were purchased from LICOR. Images were acquired using Odyssey Imaging Systems (LICOR). ImageJ software was used to perform densitometry analyses of Western blots. Results for each band were normalized to total protein levels in the same blot. GAPDH is shown as a loading control. See antibody information in Supplemental Table 9.

**Glycolysis stress test in Seahorse assay**

To study the effect of SYK inhibition in glycolysis, we profiled the glycolytic activity in cells with or without the acute treatment of SYK inhibitor (PRT062607 HCl) using Glycolysis Stress Test Kit in Seahorse XFe24 analyzer (Agilent). Briefly, a sensor cartridge (102342-100, Agilent) was hydrated in a Seahorse XF Calibrant (100840-000, Agilent) at 37°C in a non-$CO_2$ incubator overnight. On the day of measurement, cells (Nalm6, Kasumi2 and RS4;11) were collected and washed with PBS. After centrifugation, Cells were resuspended at the density of $2\times10^6$ per mL in Seahorse XF RPMI medium (pH 7.4) supplemented with 2mM glutamine without glucose and pyruvate. 100 µl cell suspension was added into each well of Seahorse XF Cell Culture Microplate (102342-100, Agilent). After centrifugation at 200 × *g* for 2 min with no brake, the plate was equilibrated for 30 min in a 37°C incubator without $CO_2$. Additional 500ul medium were added and incubated for 30 min before loading to the machine. The ECAR was measured at basal condition and sequentially after injecting control or 5µM SYKi followed by glucose, oligomycin, and 2-deoxy-D-glucose. The process measures glycolytic function by tracking extracellular acidification rate (ECAR) to assess glycolysis, glycolytic capacity, and glycolytic reserve. In addition, to confirm the effect of SYK inhibition in glycolysis sustain over time, we treated cells with SYK inhibitor for 24 hours and then measured the glycolytic function compared to untreated cells by using the Glycolysis stress test kit in Nalm6 and 697 cells following the procedures as described above.

**Signaling status and glucose dependency in cells after TKI treatment**

Cells were serum starved overnight and followed by treatment of different TKIs target S6K1 (PF-4708671) PI3K (LY294002), mTOR (rapamycin), SYK (PRT062607 HCl) as well as control for 24 hours. One million cells of each condition were harvested and proceed with CyTOF profiling. In the rest of cells, the treatments were washed out from the medium. And untreated and TKI-treated cells were cultured in the condition of glucose deprivation (GD) condition in the density of

0.5 million per mL for another 24h. Cell viability was measured by CellTiter-Glo Luminescent Cell Viability Assay (Promega) and normalized to control condition.

**Manual gating in healthy BM and primary patient samples**

Single cells were gated using Omiq software (https://www.omiq.ai/) based on event length and 191Ir/193Ir or 103Rh DNA content to filter out debris and doublets, as previously described[72]. After gating for single cells, live non-apoptotic cells were identified by gating on cleaved poly(ADP-ribose) polymerase (cPARP), cleaved caspase-3 (c-Caspase3), and 195Pt levels[68]. In PDX samples, murine cells were excluded by gating for mouse CD45 (mCD45). Platelets and erythrocytes were removed by gating on CD61 and CD235a, while T cells and myeloid cells were excluded based on CD3e, CD33, and CD16 expression. $CD38^{high}$ plasma cells were also gated out, leaving a population defined as lineage-negative blasts (Lin− B+). Unless noted otherwise, further analysis was performed on this Lin− B+ population.

**Depmap publicly available data analysis**

Depmap Batch corrected Expression Public 24Q2 data was downloaded from https://depmap.org/portal and used for the analysis to compare the batch corrected gene expression log2(TPM+1) of *UPP1*, *CDA* and *UCK1* in 1437 cell lines from 31 different cancer types [11]. The ratio of UCK1 to UPP1 expression was evaluated.

**Ribose rescue after glucose deprivation**

To rescue the effect of glucose deprivation (GD), cells were seeded in 96-well plates at 3-5 x $10^4$ cells in 200 µl of growth medium and supplemented with uridine (2mM) or ribose (10mM) under glucose deprivation condition (10% dialyzed FBS, 0mM glucose, 2mM glutamine) every 24 hours at 37 °C, 5% $CO_2$. After 48 hours, cell viability was measured by Annexin V/7AAD staining in FACS.

**The effect of DHODHi treatment at single cell level profiled by CyTOF**

Nalm16 and SJ45503 cells were seeded at the density of 0.5 million per mL in 10% FBS RPMI-1640. Cells were collected before treatment (Day0) and after BAY-2402234 treatment (100nM in Nalm16 and 500nM in SJ45503) for 3 and 6 days were profiled by CyTOF. Live cells were manually gated by excluding dead cells that are Rh103-DNA low, cleaved caspase-3 and cleaved

PARP positive. UMAP plots were generated in omiq. pS6+ live cells were further gated in UMAP plots.

### Extended Data Figures

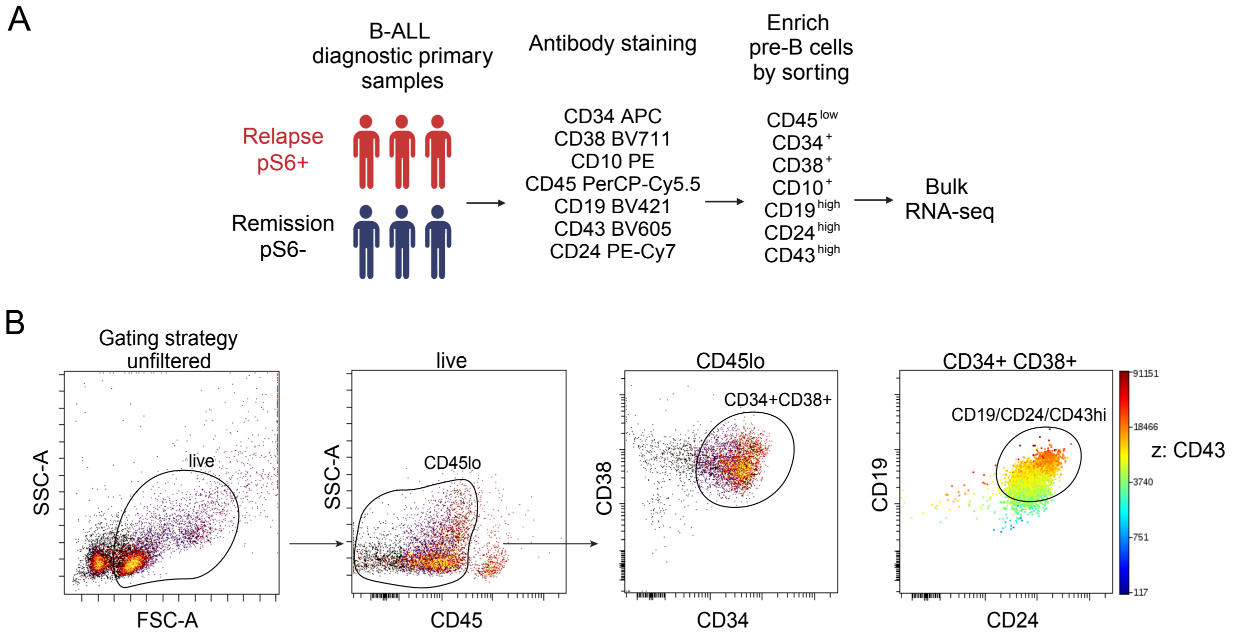


**Extended Data Fig. 1: Sorting strategy for pre-B cells from primary patient samples**

**A**, Schematic figure showing the workflow to enrich the pre-B cells from patient samples by sorting with 7-color flow cytometry staining panel. The sorted cells are processed with bulk RNA-seq.

**B**, Gating strategy of pre-B cells in primary patient samples. Pre-B cells are enriched in the population with low CD45 expression, positive for CD34, CD38 and CD10 and high expression of CD19, CD24 and CD43.

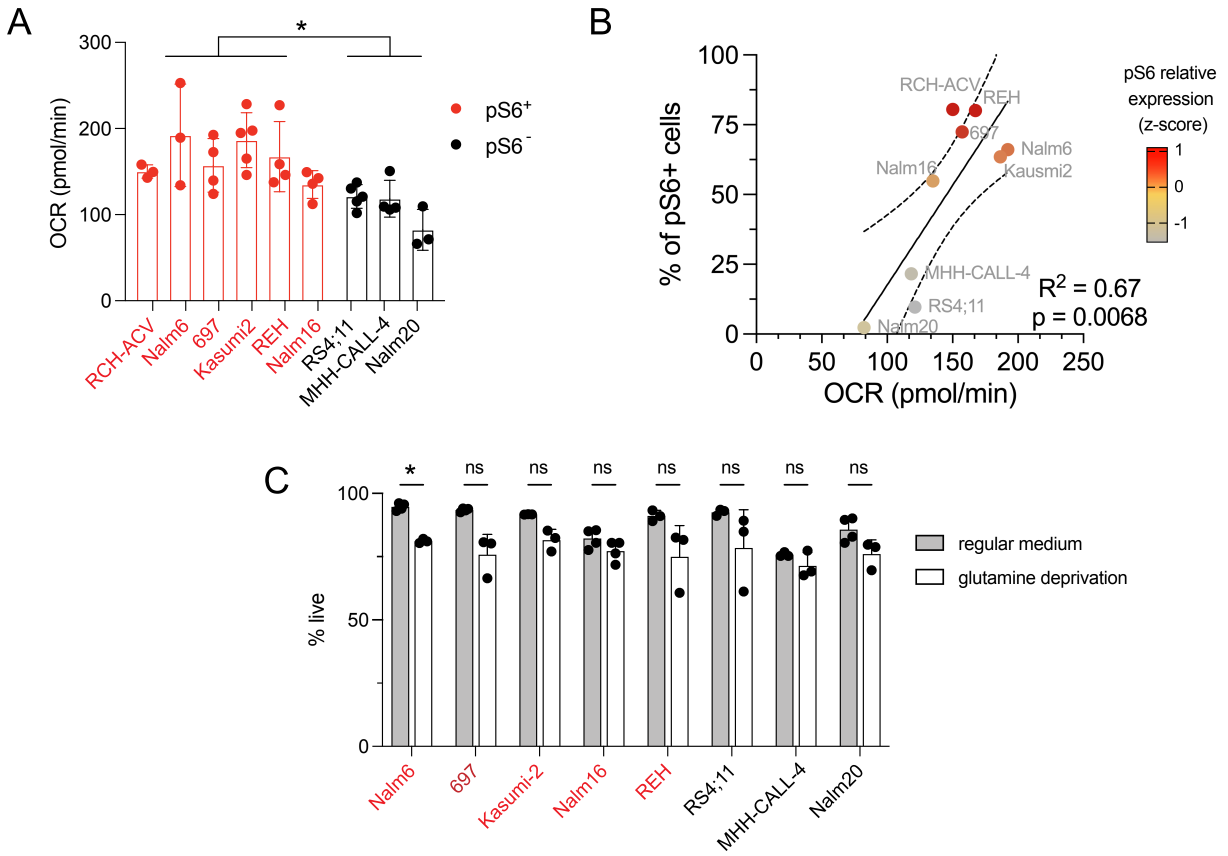


**Extended Data Fig. 2:** B-ALL cells are not dependent on glutamine for survival

**A**, Oxygen consumption rate (OCR) measured by Seahorse assay indicating mitochondrial respiration activity in pS6+ cell lines (red) compared to pS6- cell lines (black). p=0.0191

**B**, Linear regression curve demonstrating the correlation between the frequency of pS6+ cells and the OXPHOS activity (measured by OCR) in different B-ALL cell lines (n=9, p = 0.0068, $R^2$ = 0.67). Each dot represents individual cell line colored by pS6 relative expression level (z-score of arcsinh transformed mean value) measured by mass cytometry.

**C**, % of live cells in cell lines being cultured with medium with or without L-glutamine for 48 hours (n=8). Cell apoptosis is measured by annexin V and PI staining in flow cytometry. Annexin V and PI double negative cells are live cells.  All data are mean ± SD from biological three or four experiments. Statistical tests were Welch's t test (A) and multiple paired t test with correction using the Šidák-Bonferroni method. *p < 0.05, **p < 0.01, ***p < 0.001.

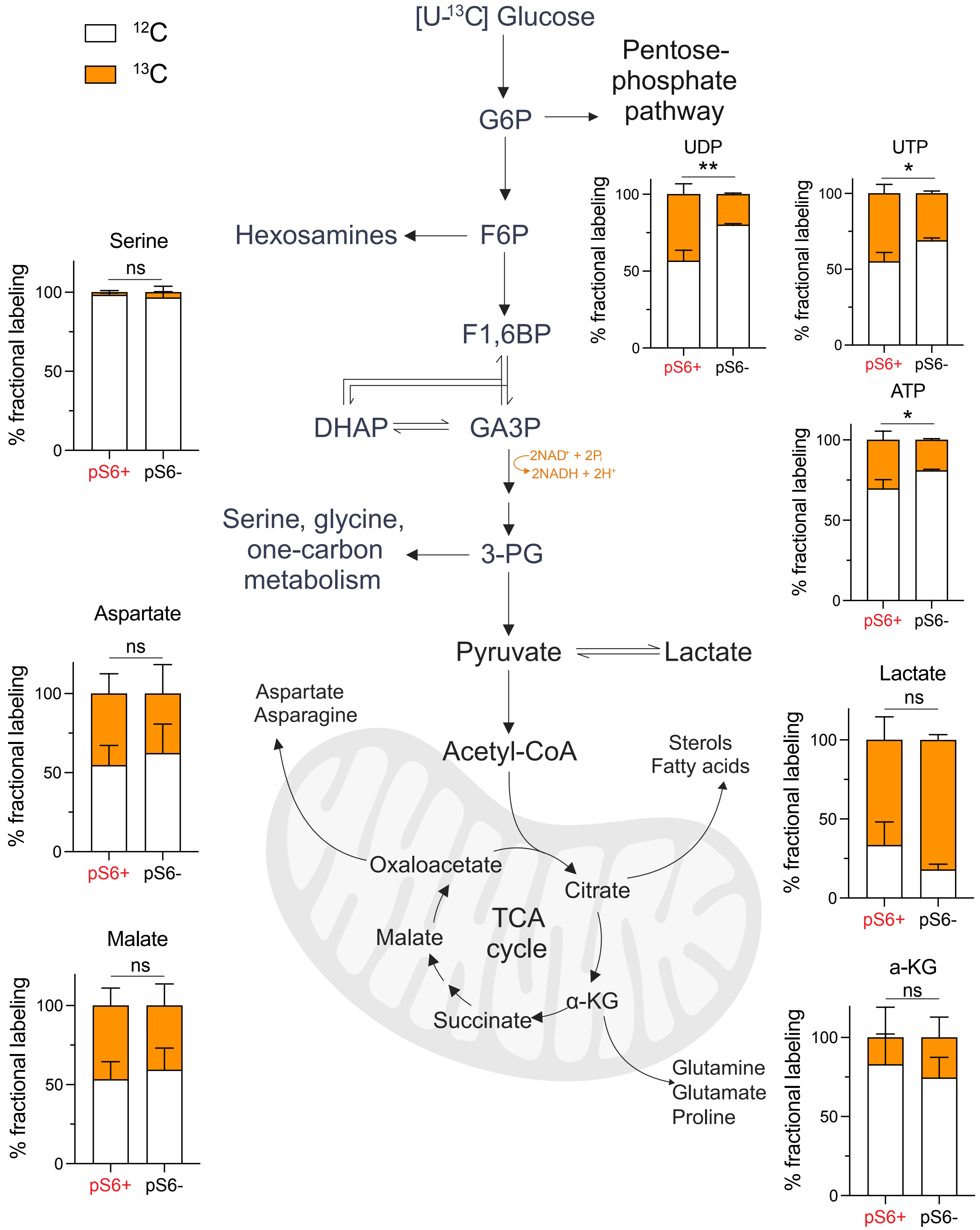


**Extended Data Fig. 3:** The fractions of $^{13}C$ labelled metabolites and unlabeled $^{12}C$ metabolites in glucose metabolism, including glycolysis, pentose phosphate pathway, serine synthesis pathway and TCA cycle in pS6+ (n=4) and pS6- (n=2) cells. Six cell lines were cultured in the condition of glutamine deprivation for 4h and fed with [U-$^{13}$C]-glucose medium without glutamine for 4h. There are significantly higher $^{13}$C labeling in uridine diphosphate (UDP, $p = 0.006$, uridine triphosphate (UTP, $p = 0.015$) and adenosine triphosphate (ATP, $p = 0.024$) in pS6+ cells compared to pS6- cells. Data in bar graph are presented as mean ± SD in triplicate. Statistical test used to compare the fractional $^{13}$C labelling between pS6+ and pS6- cells is Welch's t test. ns, not significant; *$p < 0.05$, **$p < 0.01$.

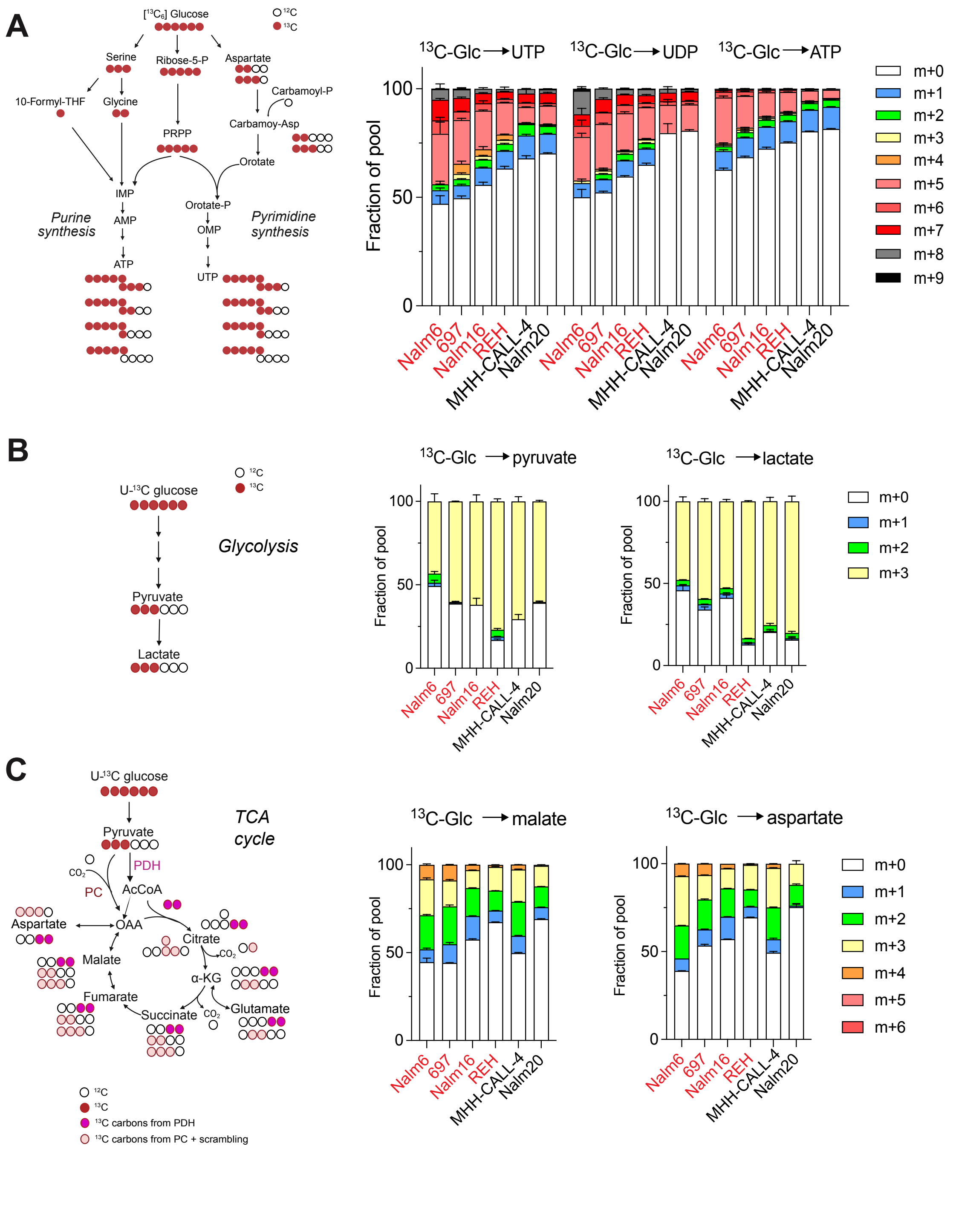


**Extended Data Fig. 4**: Isotypic tracing of $^{13}C_6$ glucose labelled metabolites in glycolysis and TCA cycle. $^{13}C_6$ glucose labeling of intermediates in pentose phosphate pathway (A), glycolysis (B) and TCA cycle (C) in B-ALL cell lines (n=6). pS6+ cells (in red): Nalm6, 697, Nalm16 and REH; pS6- cells: MHH-CALL-4, Nalm20. Cells were cultured in the condition of glutamine deprivation for

8h and fed with [U-$^{13}$C]-glucose medium without glutamine for 4h. Data are presented as mean ± SD in triplicates. Data in this figure was after cell lysate normalization.

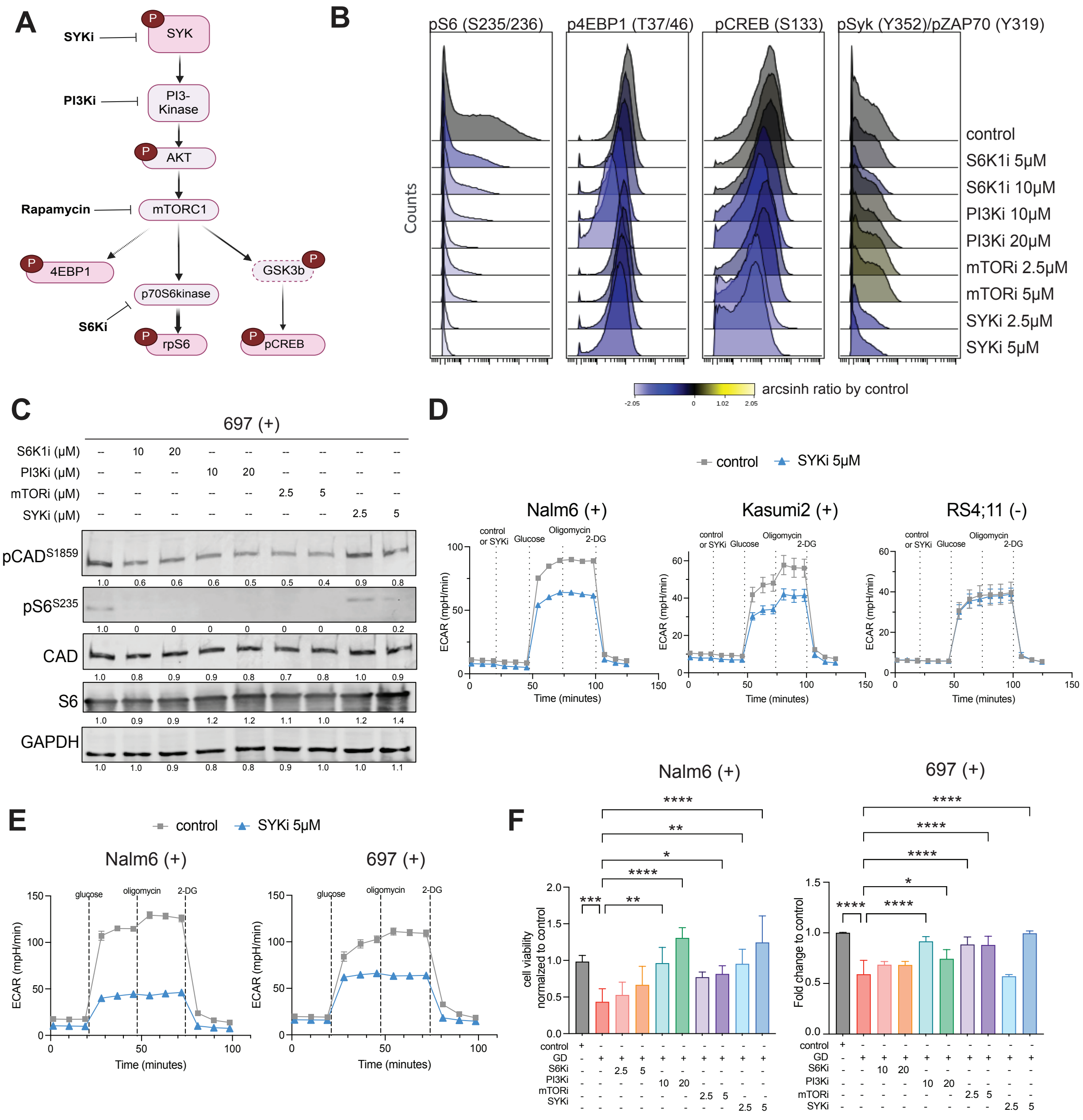

**Extended Data Fig. 5**: Glucose-dependent uridine synthesis is driven by pS6 signaling regulated by mTOR/PI3K axis.

A) Schematic figure of SYK/PI3K/mTOR signal axis.

B) Histograms to show phosphorylation of ribosome protein S6 (S235/236), 4EBP1 (T37/46), pCREB (S133) and pSYK (Y352) after treatment with tyrosine kinase inhibitors to target S6K1 (PF-4708671: 10, 20μM) PI3K (LY294002: 10, 20μM), mTOR (rapamycin: 2.5, 5μM), SYK (PRT062607 HCl: 2.5, 5μM) for 24 hours. Protein profile was measured by CyTOF and normalized to control condition. S6K1i inhibits pS6; PI3Ki inhibits pS6 and p4EBP1; mTORi inhibits pS6; and SYKi inhibits pS6, p4EBP1, pCREB and pSYK.

C) Western blot analysis showing the effect of tyrosine kinase inhibitions by small inhibitors in phosphorylation of S6 (S235/236) and CAD (S1859) in cell line 697. After serum starvation overnight, cells were treated with different tyrosine kinase inhibitors to target S6K1 (PF-4708671: 10, 20μM) PI3K (LY294002: 10, 20μM), mTOR (rapamycin: 2.5, 5μM), SYK (PRT062607 HCl: 2.5, 5μM) for 24 hours, followed by cytoplasmic protein extraction and analysis by western blot.

D) Extracellular acidification rate (ECAR) profile in cells with or without the acute treatment of SYKi. ECAR in the cells were measured by Seahorse analysis in different time points with sequential injections of control or SYKi followed by glucose, Oligomycin and 2-DG in pS6+ cells (Nalm6, Kasumi-2) and pS6- cells (RS4;11), respectively.

E) ECAR profile in cells with or without 24-hour SYKi treatment. Cell lines (Nalm6 and 697) were treated in control and SYKi (5 μM) for 24 hours and followed by the glycolysis stress test in Seahorse.

F) Bar graphs to show the effect of tyrosine kinase inhibitions in cell viability under glucose deprivation condition. Cells were serum starved overnight and followed by treatment with different tyrosine kinase inhibitors in the condition of glucose deprivation (GD) for 24h. Cell viability was measured by cell titer glo assay and normalized to control condition.

All data in bar graph are mean ± SD. Statistical test used is one-way ANOVA followed by Dunnett's multiple comparison test (C). $^{*}p < 0.05$, $^{**}p < 0.01$, $^{***}p < 0.001$, $^{****}p < 0.0001$

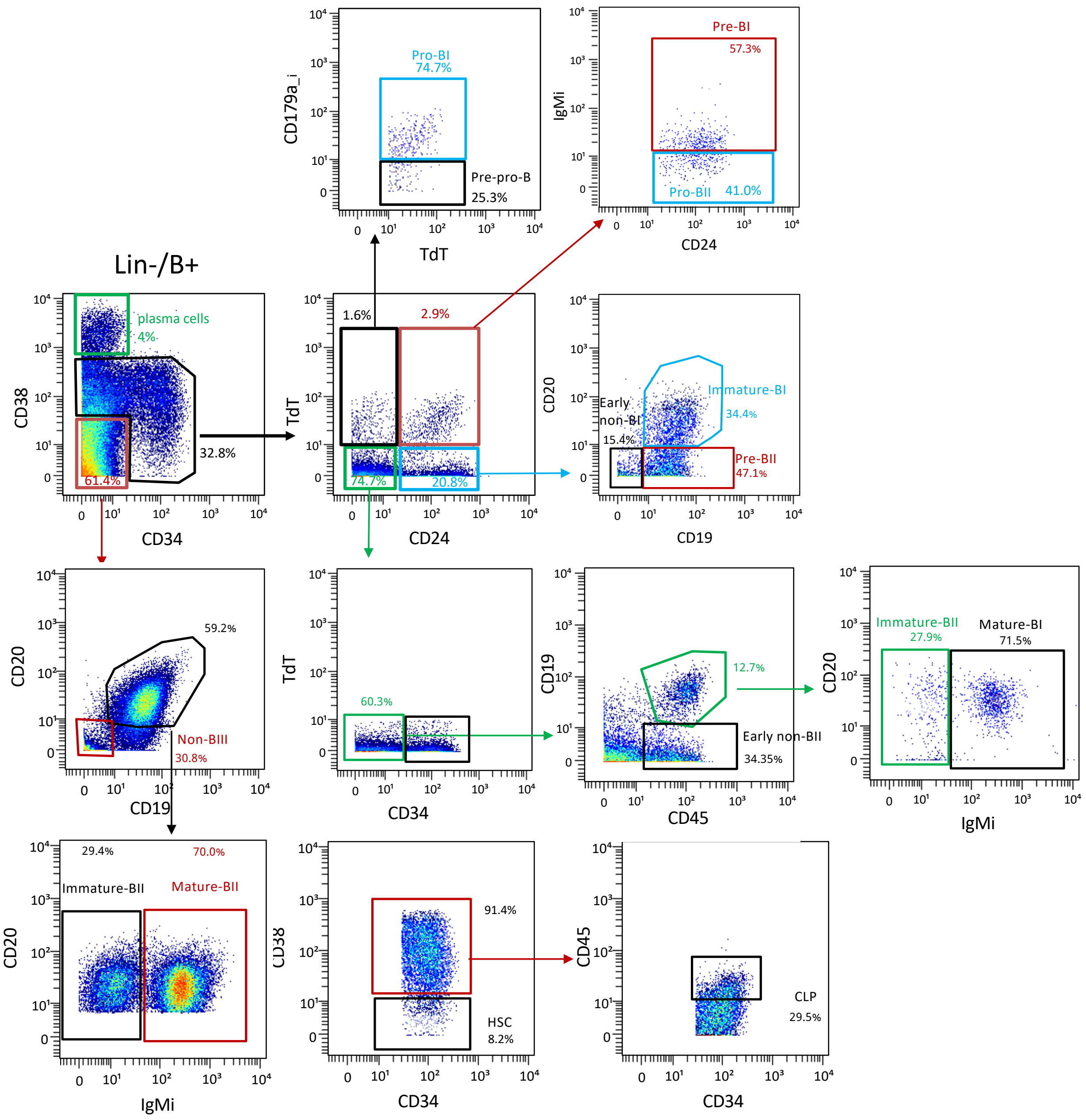


**Extended Data Fig. 6**: Manual gating strategy for healthy BM sample. The actual gating from the lineage negative blast starts with the second-row left plot (CD34xCD38). Percentages are all relative to the individual plots. The resulting 15 subpopulations used in the developmental classifier are labeled.

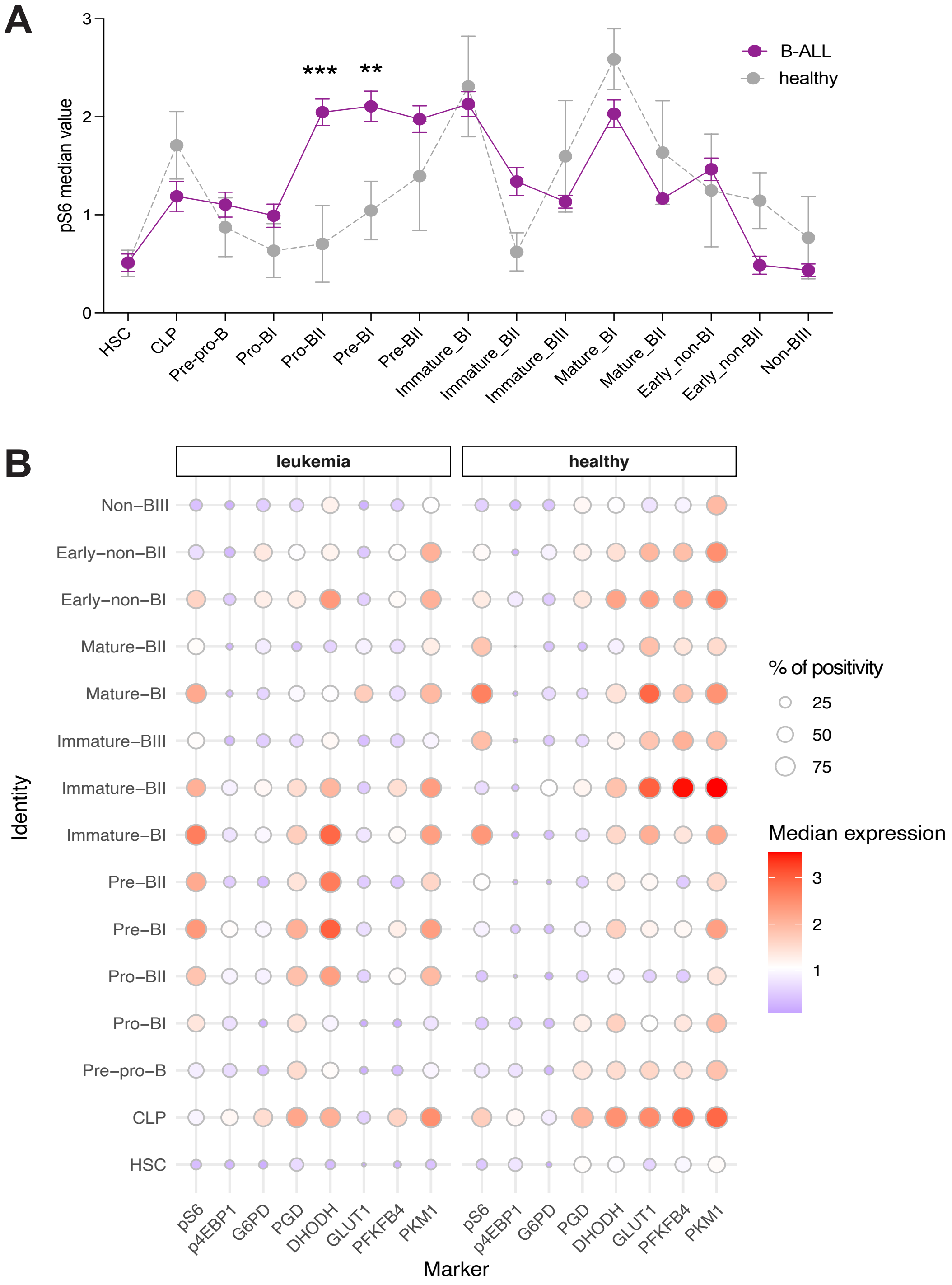


**Supplement Figure 7**: pS6 signaling strength and metabolic protein expression in different B cell developmental stages in primary B-ALL patient samples and healthy bone marrows.

A) Median (arcsinh transformed) value of pS6 (S235/236) in classified cell populations in B-ALL (purple) compared to healthy bone marrows (gray). B-ALL primary patient samples (n=31) as well as healthy bone marrows (n=5) were profiled by CyTOF. Protein profiles were used for developmental classification. B) Median expression (arcsinh transformed) and percent positivity of proteins in classified cell populations in B-ALL samples (n=31) and healthy BMs (n=5), respectively. B-ALL primary patient samples (n=31) as well as healthy bone marrows (n=5) were profiled by CyTOF. Protein profiles were used for developmental classification.

Data was shown in mean ± SEM. Statistical test used was 2way-ANOVA test followed by Šidák test. *$p < 0.05$, **$p < 0.01$, ***$p < 0.001$.

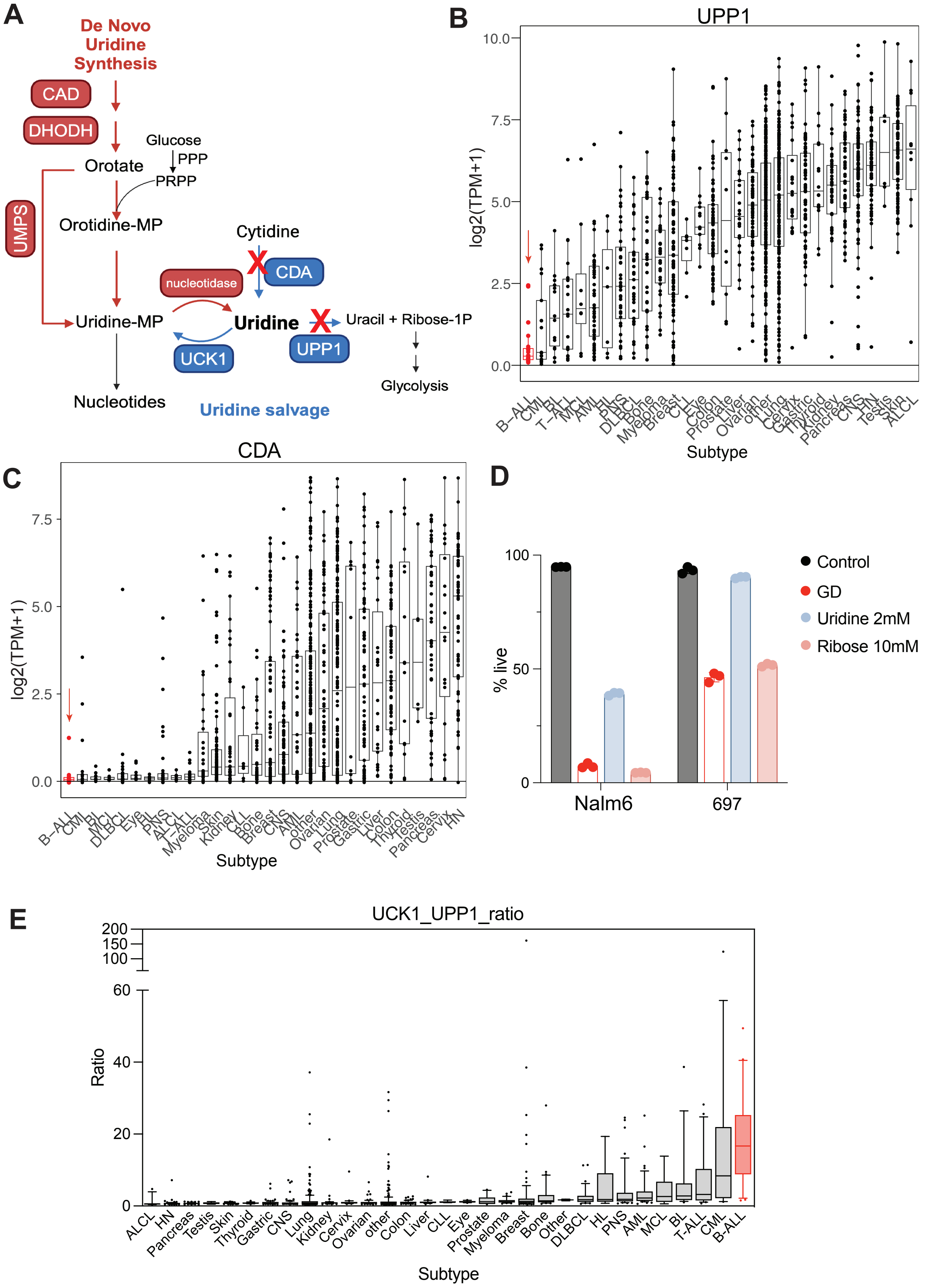
A
De Novo Uridine Synthesis
CAD
DHODH
Glucose
PPP
Orotate
PRPP
UMPS
Orotidine-MP
Cytidine
CDA
nucleotidase
Uridine-MP
Uridine
Uracil + Ribose-1P
UPP1
UCK1
Nucleotides
Glycolysis
Uridine salvage
B
UPP1
log2(TPM+1)
Subtype
C
CDA
log2(TPM+1)
Subtype
D
% live
Nalm6
697
Control
GD
Uridine 2mM
Ribose 10mM
E
UCK1_UPP1_ratio
Ratio
Subtype

**Extended Data Fig. 8**: Uridine salvage through UPP1 and CDA is not active in B-ALL.

A. Schematic figure to show that uridine can be synthesized from orotate and ribose-5P in de novo synthesis pathway indicated as red arrows and salvaged from cytidine indicated as blue arrows. Other studies have shown that uridine can be converted to uracil and ribose-1P to fuel glycolysis.
B. Gene expression of UPP1 (encoding uridine phosphorylase, a pyrimidine salvage enzyme that catalyzes the reversible phosphorylation of uridine to uracil and ribose-1-phosphate) in 1437 cell lines from 31 different cancer types in Depmap RNA-seq data (Batch Corrected Expression Public 24Q2). The red arrow indicates B-ALL cell lines have the lowest expression of UPP1 crossing all cancer types.
C. Gene expression of CDA (encoding cytidine deaminase, an enzyme involved in uridine salvage pathway that converts cytidine to uridine) in Depmap RNA-seq data (Batch Corrected Expression Public 24Q2). The red arrow indicates B-ALL cell lines have the lowest expression of CDA crossing all cancer types.
D. Cell viability after glucose deprivation for 48 hours with and without rescuing by uridine (2mM) or ribose (10mM). pS6+ cells are rescued by uridine or ribose supplementations in the face of glucose deprivation (GD). Cell apoptosis is measured by annexin V and PI staining in flow cytometry. Annexin V and PI double negative cells are live cells.
E. Ratio of gene expression of UCK1 (uridine cytidine kinase 1) to UPP1 in 1437 cell lines from 31 different cancer types. B-ALL cell lines ranked highest (red).

B-ALL, B-cell lymphoblastic leukemia; CML, chronic myeloid leukemia; BL, Burkitt lymphoma; AML, acute myeloid leukemia; MCL, mantle cell lymphoma; HL, Hodgkin lymphoma; PNS, Peripheral Nervous System; CNS, Central Nervous System; ALCL, Anaplastic Large-Cell Lymphoma; T-ALL, T cell lymphoblastic leukemia/lymphoma; HN, Head and Neck.

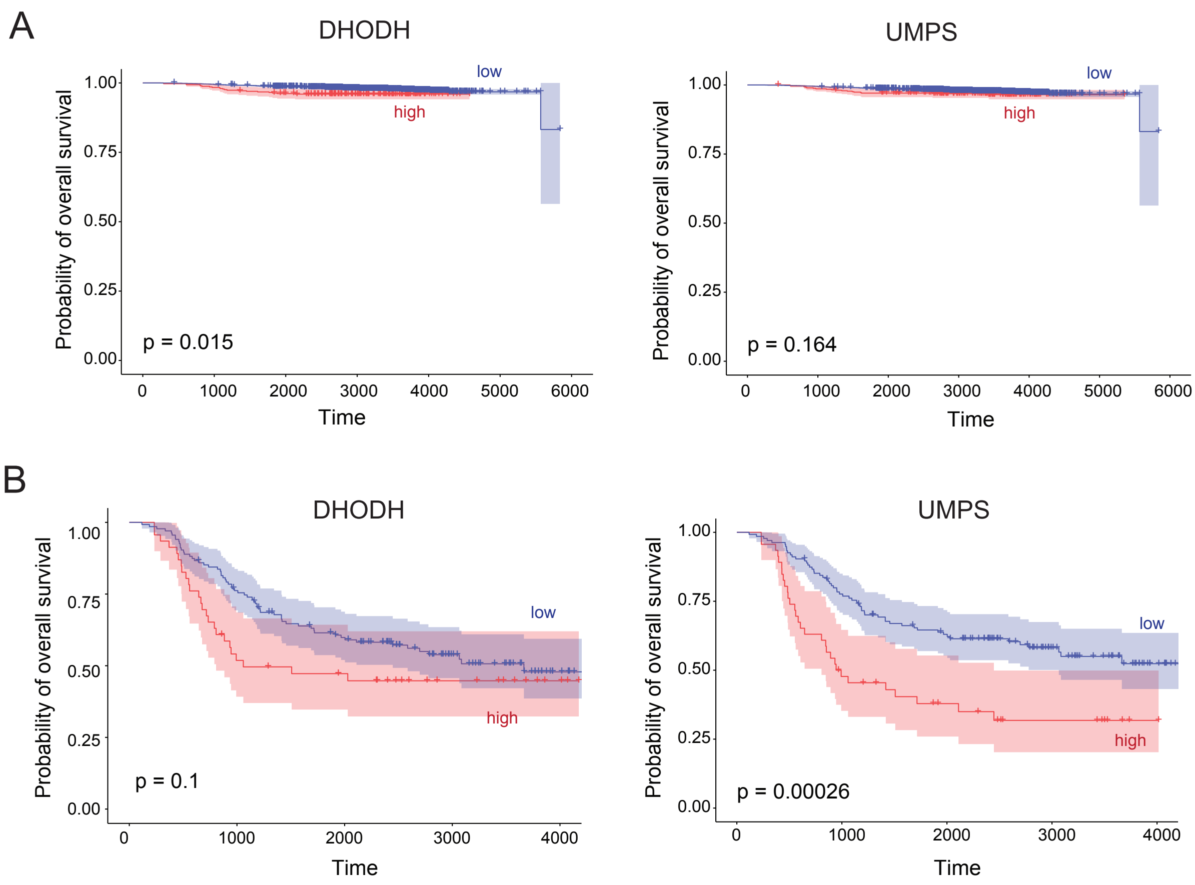


**Extended Data Fig. 9**: Overall survival of DHODH and UMPS in MP2PRT and TARGET datasets.

A) Weighted Kaplan-Meier curves comparing probability of overall survival in groups with different ranking of DHODH and UMPS expression in MP2PRT dataset. N= 147 in the highest 10-percentile group; n=1,318 in the lowest 90-percentile group. MP2PRT, Molecular Profiling to Predict Responses to Therapy. Significance values from weighted Cox regression test results are shown.

B) Kaplan-Meier curves comparing probability of overall survival in patients with different ranking of DHODH and UMPS expression in NCI TARGET dataset. n = 46 in the highest 25-percentile group; n = 135 in the lowest 75-percentile group. The Cox regression test was used for the survival curves analyses.

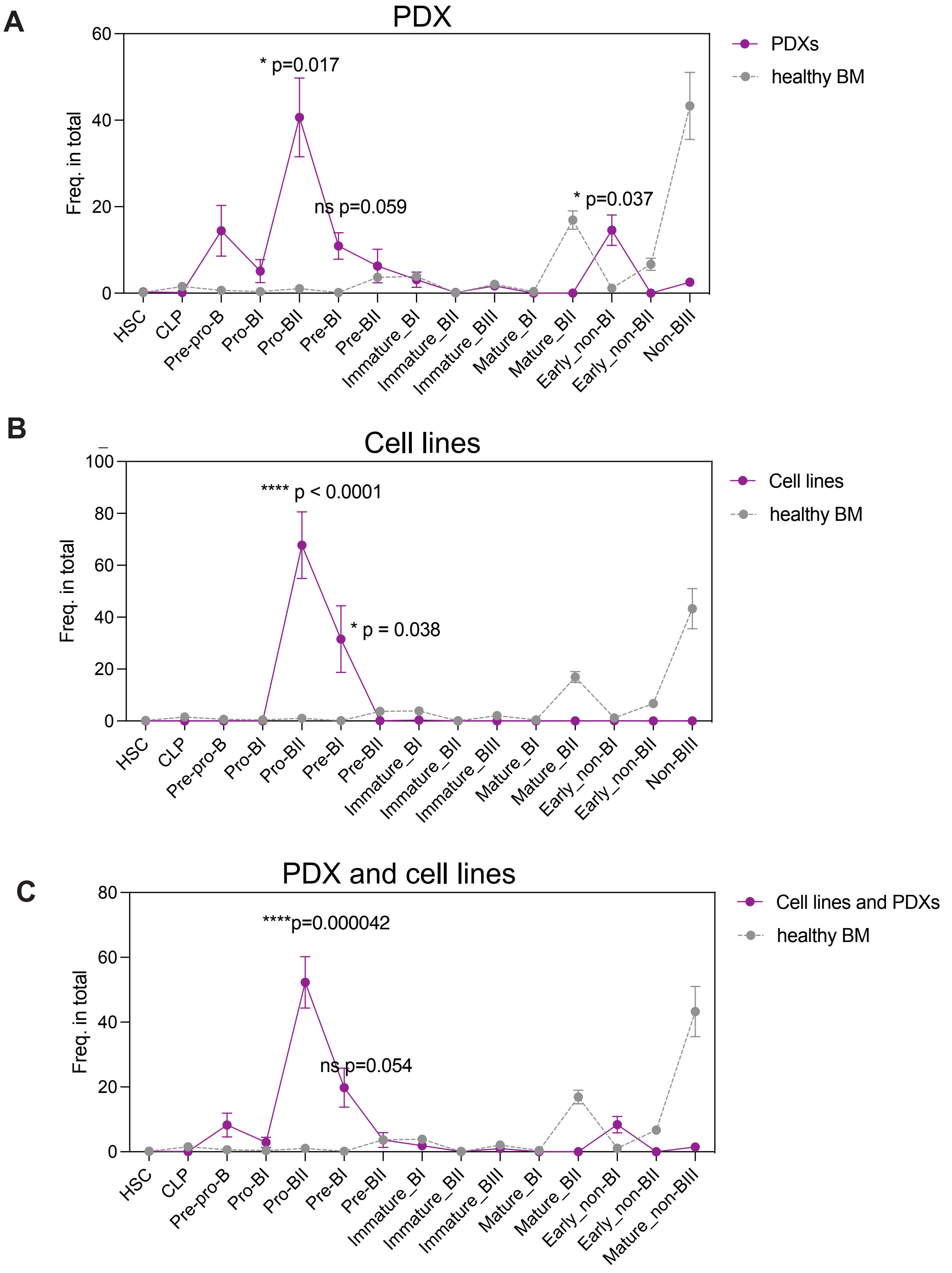


**Extended Data Fig. 10**: Enrichment of Pro-BII and Pre-BI as demonstrated by developmental classification in PDX samples (A, n=12) and cell lines (B, n=9). All combined is shown in C.

Protein profiles were measured by CyTOF. Lineage-B+ cells were manually gated in omiq and exported for developmental classification in R. Data are shown in mean ± SEM. Statistical test used to compare the frequency of subpopulations in total between samples (purple line) and healthy BM (gray line, n=2) was multiple unpair t test and adjusted by Holm-Šidák method. ns, not significant, *p < 0.05, **p < 0.01, ***p < 0.001.

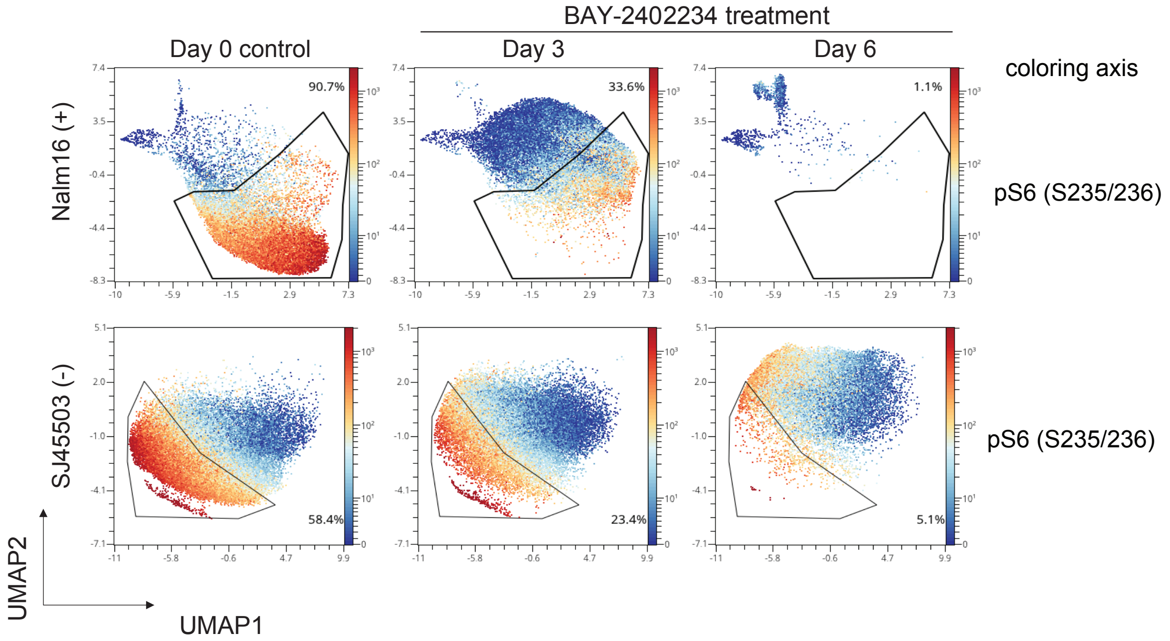


**Extended Data Fig. 11**: DHODH inhibition selectively targets pS6+ cells over time.

Expression of pS6 and frequency of pS6+ cells after DHODH inhibition in NALM 16 cells (pS6+, first row) and PDX SJ45503 (pS6-, second row). Nalm16 and SJPDX cells collected before (Day 0) and after BAY-2402234 treatment for 3 and 6 days were profiled by CyTOF. We first gated on live cells by excluding dead cells that are Rh103-DNA low, cleaved caspase-3 and cleaved PARP positive and generated UMAP plots in omiq. pS6+ live cells were further gated in UMAP plots. The coloring axis in UMAP is pS6 (S235/236).

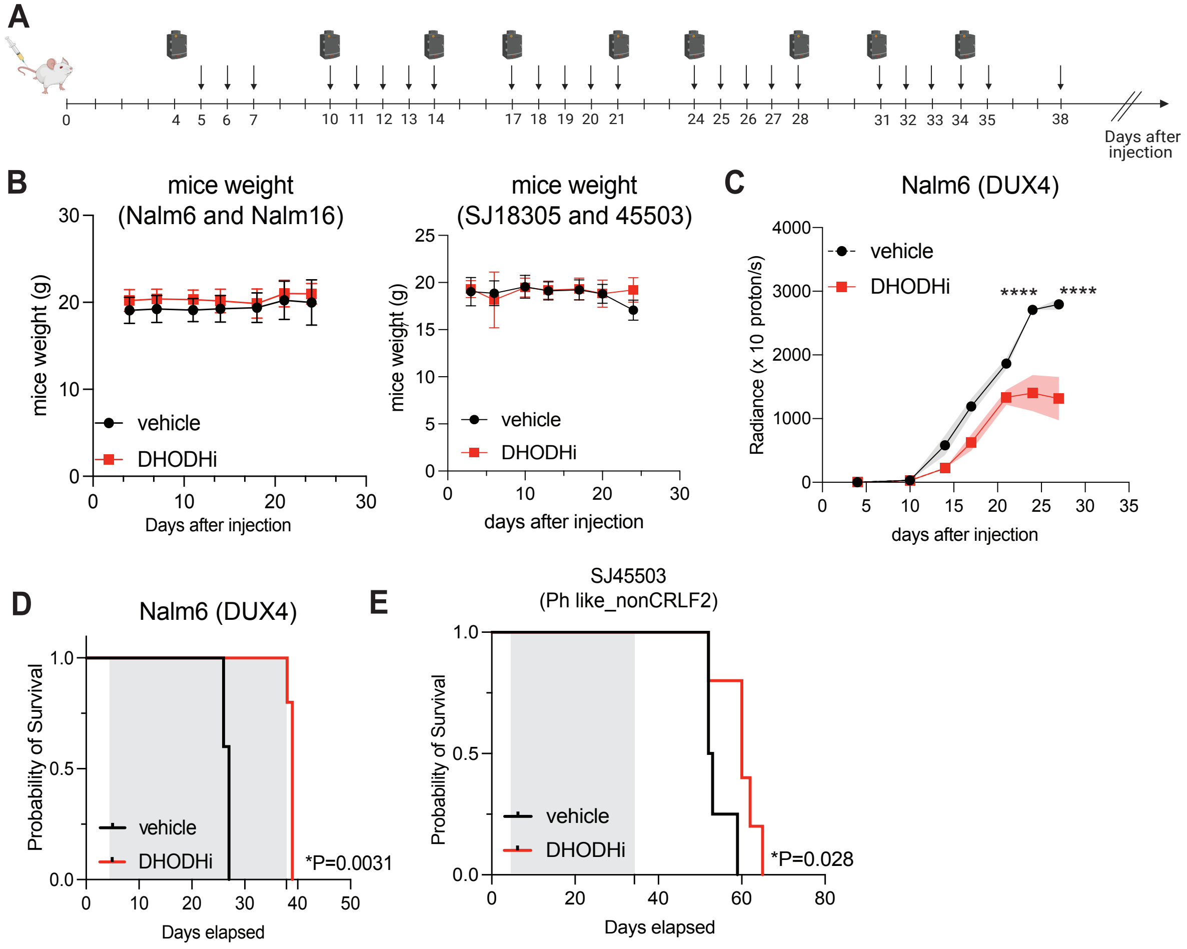
A
0
4 5 6 7
10 11 12 13 14
17 18 19 20 21
24 25 26 27 28
31 32 33 34 35
38
Days after injection
B
mice weight (Nalm6 and Nalm16)
mice weight (g)
Days after injection
vehicle
DHODHi
mice weight (SJ18305 and 45503)
mice weight (g)
days after injection
vehicle
DHODHi
C
Nalm6 (DUX4)
Radiance (x 10 proton/s)
days after injection
vehicle
DHODHi
****
****
D
Nalm6 (DUX4)
Probability of Survival
Days elapsed
vehicle
DHODHi
*P=0.0031
E
SJ45503
(Ph like_nonCRLF2)
Probability of Survival
Days elapsed
vehicle
DHODHi
*P=0.028

**Extended Data Fig. 12**: DHODH inhibition is preferentially effective in xenografts with pS6 activation.

A) *In vivo* dosing schedule: 2 x $10^5$ cells were tail vein injected to each NSG mice. Nalm6 and Nalm16 xenografts were treated daily with 5mg/kg DHODHi (BAY-2402234) for 24 dosing days (skip weekends). The treatment stopped at $38^{th}$ day after iv injection. BLI was performed twice a week.
B) Weight was measured twice a week in Nalm6/Nalm16 xenografts (left panel) and SJ18305/SJ44503 xenografts (right panel) for both vehicle and DHODHi treated mice.
C) Curves to show luciferase labeled tumor growth in Nalm6 xenografts by bioluminescence in DHODHi (red curve) treated and vehicle (black curve) mice.
D) Survival of Nalm6 xenografts treated by DHODHi (red curve) and vehicle mice (black curve).
E) Survival of SJ45503 PDX xenografts treated by DHODHi (red curve) and vehicle mice (black curve).

Data in curves (B) are mean ± SD. Data in curves (C and E) are mean ± SEM represented as dots and area fill within error bands and a two-way ANOVA mixed model followed by Sidak's test for multiple comparisons was used. Log-rank test was used in Kaplan Meier curves (D and F). ns, not significant, *$p < 0.05$, **$p < 0.01$, ***$p < 0.001$, ****$p < 0.0001$.

**Supplemental Information**

**Supplemental Table 1**. Primary patient samples from Good et al. Nature Med 2018 for transcriptomic sequencing of sorted pre-B cells.

**Supplemental Table 2**. Flow cytometry antibodies for sorting pre-B cells from primary patient samples

**Supplemental Table 3**. GSEA pathway enrichment from diagnostic pS6+ pre-B cells from patients who would go on to relapse compared to pS6- pre-B cells from patients in continuous remission. Hallmark database was used in GSEA analysis. FDR $< 0.25$.

**Supplemental Table 4**. CYTOF panel used for healthy bone marrows, cell lines and primary samples.

**Supplemental Table 5**. Information for patient samples utilized from Bass Center Tissue Biobank at Stanford.

**Supplemental Table 6**. GSEA pathway enrichment analysis in patients who would go on relapse compared to patients who are in remission in MP2PRT and TARGET datasets.

**Supplemental Table 7**. PDX sample information.

**Supplemental Table 8**. Nutrient supplements used in rescue experiments.

**Supplemental Table 9**. Western blot antibodies
